# A molecular brake that modulates spliceosome pausing at detained introns contributes to neurodegeneration

**DOI:** 10.1101/2022.01.11.475943

**Authors:** Dawei Meng, Qian Zheng, Xue Zhang, Li Luo, Yichang Jia

**Author notes:** Corresponding author, Please address correspondence to: Yichang Jia, Ph. D., School of Medicine, Medical Science Building, Room D204, Tsinghua University, Beijing, 100084, P. R. China.

## Abstract

Emerging evidence suggests that intron-detaining transcripts (IDTs) are a nucleus-detained and polyadenylated mRNA pool for cell to quickly and effectively respond to environmental stimuli and stress. However, the underlying mechanisms of detained intron (DI) splicing are still largely unknown. Here, we suggest that post-transcriptional DI splicing is paused at B^act^ state, an active spliceosome but not catalytically primed, which depends on SNIP1 (Smad Nuclear Interacting Protein 1) and RNPS1 (a serine-rich RNA binding protein) interaction. RNPS1 and B^act^ component preferentially dock at DIs and the RNPS1 docking is sufficient to trigger spliceosome pausing. Haploinsufficiency of *Snip1* attenuates neurodegeneration and globally rescues IDT accumulation caused by a previously reported mutant U2 snRNA, a basal spliceosomal component. *Snip1* conditional knockout in cerebellum decreases DI splicing efficiency and causes neurodegeneration. Therefore, we suggest that SNIP1 and RNPS1 form a molecular brake for the spliceosome pausing, and that its misregulation contributes to neurodegeneration.

## Introduction

Introns are spliced from eukaryotic messenger RNA precursors (pre-mRNA) by the spliceosome via two transesterification reactions - branching and exon ligation (*1*). During these reactions, the spliceosome undergoes structural and compositional dynamics (*2–4*). Firstly, the 5’ splice site (SS), branch site (BS), and 3’SS of an intron are recognized by the U1 small nuclear RNA (snRNA), SF1, and U2AF, respectively (E complex). Then, U2 small nuclear ribonucleoprotein particles (snRNPs) are recruited by the E complex to the BS (A complex), which then binds to U4/U6.U5 tri-snRNP to form the fully assembled and pre-catalytic spliceosome (B complex). The resulting B complex is empowered by an ATPase/helicase Brr2, remodeling to the active spliceosome (B^act^). Through additional remodeling by the ATPase/helicase Prp2, B^act^ matures into a catalytically activated spliceosome (B*), in which the branching reaction occurs. During the B-to-B^act^ and B^act^-to-B* transitions, a number of proteins are loaded/unloaded into the spliceosome, including more than 10 proteins recruited into early B^act^ and release of SF3a, SF3b, and pre- mRNA REtention and Splicing (RES) complexes from B* (*5–8*). The resulting catalytic step I spliceosome (C complex) is remodeled by the ATPase/helicase Prp16 into a step II catalytically activated spliceosome (C* complex), in which the exon ligation reaction occurs.

In contrast to constitutive splicing, > 90% of human multiexon genes undergo alternative splicing (AS) (*9, 10*), which contributes to proteomic diversity (*11, 12*). Accuracy in the recognition of reactive splice sites must be compromised by flexibility in splice site choice during AS. As one of the major categories of AS, intron retention (IR) was originally thought to be nonproductive for protein production, because it often introduces a premature termination codon (PTC) into the transcript, which is subsequently targeted for degradation by nonsense-mediated decay (NMD), a cytoplasmic mRNA surveillance mechanism (*13, 14*). However, Boutz et al. identified a group of intron-detaining transcripts (IDTs) in human and mouse cells that are polyadenylated, detained in the nucleus, and immune to NMD, and termed these incompletely-spliced introns as detained introns (DIs) (*15*). Nucleus DIs have also been documented by others and their splicing and subsequent mRNA export to the cytoplasm for protein translation has been associated with specific stimuli and stress (*16–23*). Given that gene size in human is large (∼27 kb in average) but RNA transcription rate is slow (∼2-4kb/minute) (*24–27*), post-transcriptional DI splicing is a quick and effective way for cells to adapt environment changes or stress. However, it remains unclear whether spliceosome is indeed paused at DIs, which catalytic step the spliceosome is paused at, what the molecular mechanisms underlie the spliceosome pausing, and what are the biological consequences of misregulation of this process.

Here, we disclose that more than one third of cerebellum-expressed genes transcribe IDTs and ∼90% of them only contain 1-2 intron(s). Using mouse forward genetics and gene knockout (KO), we demonstrate that haploinsufficiency of *Snip1* (Smad Nuclear Interacting Protein 1) rescues IDT accumulation and neurodegeneration caused by a previously reported mutant U2 snRNA (*28*). SNIP1 interacts with protein components found in B^act^ spliceosome and protein components in peripheral exon junction complex (EJC), like RNPS1 (RNA binding protein with serine rich domain 1). Like *Snip1*, knockdown of *Rnps1* rescue the DI accumulation while its overexpression is sufficient to trigger spliceosome pausing at DIs. B^act^ component and RNPS1 preferentially deposit at DIs and their surrounding sequences. Both RNPS1 docking at DIs and interaction between SNIP1 and RNPS1 are required for spliceosome pausing at DIs. *Snip1* conditional KO in cerebellum reduces DI splicing efficiency and leads to IDT accumulation and neurodegeneration. Therefore, we suggest that SNIP1 and RNPS1 function as a molecular brake for spliceosome pausing at highly regulated DIs, and that misregulation of this process contributes to the pathogenesis of neurodegeneration.

## Results

### A mouse forward genetic screening identifies *Snip1* as a modifier for *NMF291* phenotypes

To understand how the previously reported mutant U2 (*28*) leads to global RNA splicing abnormalities and massive cerebellar granule cell loss, we established an ENU-induced mutagenesis screening for dominant modifier(s) that rescue the *NMF291*^-/-^ phenotypes (Fig. S1). A modifier (*Snip1*^M/+^) partially rescued *NMF291*^-/-^ ataxia in a dominant manner (Fig. 1A and Movie 1). One of the modifier candidates in the family was a G to A substitution that alters the 5’ splice site (5’SS) GT of *Snip1* exon 2 to AT (Fig. 1B), which completely segregated with the rescue in the *NMF291*^-/-^ mice (Table S1). Because the ENU mutation disrupts the 5’SS, we generated a *Snip1* KO mouse line by Crispr- Cas9 to examine whether *Snip1*^M/+^ rescues *NMF291*^-/-^ phenotypes through a loss-of-function mechanism (Fig. 1C). Heterozygous *Snip1* KO (*Snip1*^-/+^) rescued *NMF291*^-/-^ ataxia and significantly extended *NMF291*^-/-^ life span, comparable to the extent of *Snip1*^M/+^ (Fig. 1D). In addition, *Snip1*^-/+^ partially rescued neuron loss in the *NMF291*^-/-^ cerebellum (Fig. 1E), although *Snip1*^-/+^ itself did not show cerebellar neuron loss. We failed to harvest a homozygous *Snip1* mutant mouse for both ENU-induced mutation and Crispr-Cas9- generated KO (Table S2), indicating that *Snip1* is an essential gene and ENU-induced mutation is possibly a null allele. Therefore, we conclude that haploinsufficiency of *Snip1* rescues neurodegenerative phenotypes shown in the *NMF291*^-/-^ mutant mouse.

**Figure 1.**
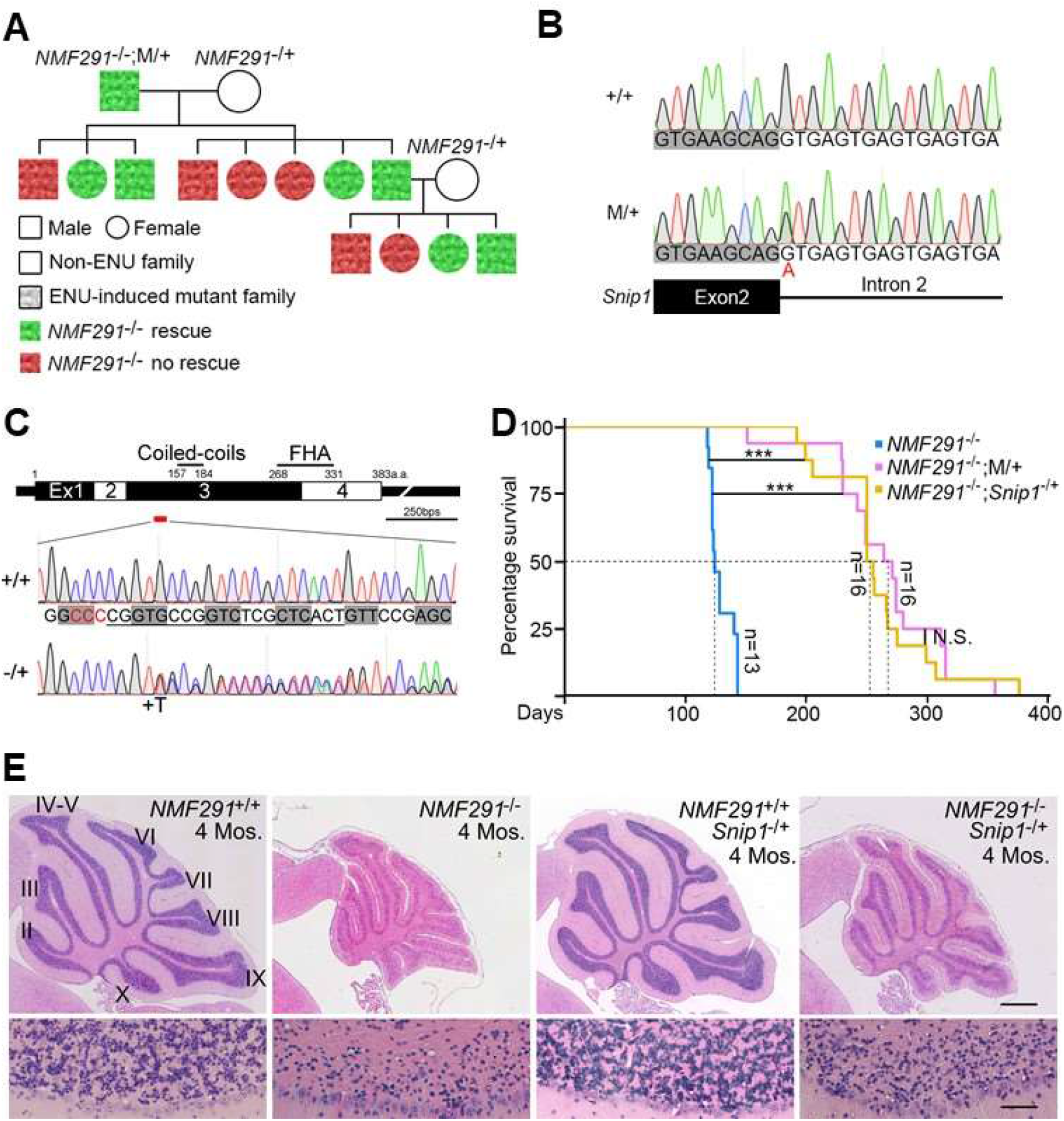
A forward genetic screening identifies *Snip1* as a modifier for *NMF291* phenotypes. (A) A *NMF291* modifier family pedigree. An ENU-induced mutant G1 male (*NMF291*^-/-^;M/+) carrying less ataxia phenotype (rescued, in green) was bred to an ENU-untreated *NMF291*^-/+^ (in white) female. The resulting *NMF291*^-/-^ mice were included for our phenotyping. The modifier is inherited in a dominant manner. (B) The ENU-induced modifier candidate (M/+) hits the 5’SS (from GT to AT) of *Snip1* exon2 (ENSMUST00000052183.6). (C) Generation of *Snip1* KO mouse (*Snip1*^-/+^) by Crispr/Cas9. *Snip1* contains 4 protein-coding exons and two functional domains. Coiled-coils, 157-184 a.a.; FHA (forkhead-associated domain), 268-331 a.a.. The Cas9 editing site was labeled as a red bar. A nucleotide insertion (+T) in *Snip1* exon3 causes out-of-frame KO. PAM site was labeled in red and gRNA sequences were underlined. Codons were labeled with gray rectangles. (D) The survival curves for the indicated genotypes (\*\*\**p*<0.001, N.S., no statistical significance, Log-rank test). Both the ENU-induced mutation (M/+) and Crispr/Cas9-generated *Snip1* KO (*Snip1*^-/+^) extended the *NMF291*^-/-^ mutant lifespans. (E) Hematoxylin and eosin-stained cerebellar sagittal sections of the indicated genotypes at 4 months (4 Mos.) of age. Lower panels, the coresponding high magnification images (lobule II). Scale bar, upper 500 μm; lower, 50 μm. See also Figure S1, Movie 1, and Table S1 and Table S2.

### Haploinsufficiency of *Snip1* globally rescues IRs in *NMF291* mutant cerebellum

One of pathological features in *NMF291* mutant cerebellum is severe IRs (*28*). To quantitatively and globally interrogate whether *Snip1*^-/+^ is able to rescue the IRs, we performed cerebellar RNA-seq with poly(A) selection in wildtype (+/+), *Snip1*^-/+^, *NMF291*^-/-^, and *NMF291*^-/-^;*Snip1*^-/+^ mice at one month of age, when the expression of mutant U2 snRNAs starts to be upregulated and IRs become severe (*28*). We employed intron retention index (IRI) to represent the IR levels, which was calculated by the ratio of intronic reads normalized to flanking exonic reads (*28–30*). Consistent with a previous report (*28*), the IRI ratio of *NMF291*^-/-^ to +/+ showed more IRs in the *NMF291*^-/-^ cerebella (Figs. 2A and S2A-S2C), regardless of whether considering high (FC > 1.2 and *p*adj < 0.1) or low (FC >1.2 and *p*adj ≥ 0.1) confidence IR events. As reported before (*28*), about half of high confidence IRs were small introns (intron length < 150bps). The comparative IRI ratio of *Snip1*^-/+^ to +/+ showed a slight depletion of IRs in *Snip1*^-/+^ cerebella (Fig. 2B). Pairwise comparison between *NMF291*^-/-^ and *NMF291*^-/-^;*Snip1*^-/+^ indicated that the majority of IRs was rescued by *Snip1*^-/+^ (Figs. 2C and S2A-S2C). Indeed, the IRI ratio of *NMF291*^-/-^;*Snip1*^-/+^ to +/+ revealed even fewer IRs shown in *NMF291*^-/-^;*Snip1*^-/+^ cerebella than that of +/+ (Fig. 2D). Among these high confident but not rescued IRs (493 shown in Fig. 2D), 74.0% are small introns with a much higher IRI ratio (median = 8.2) than that of the rest (median = 4.5), suggesting that these IRs are insensitive to haploinsufficiency of *Snip1*.

**Figure 2.**
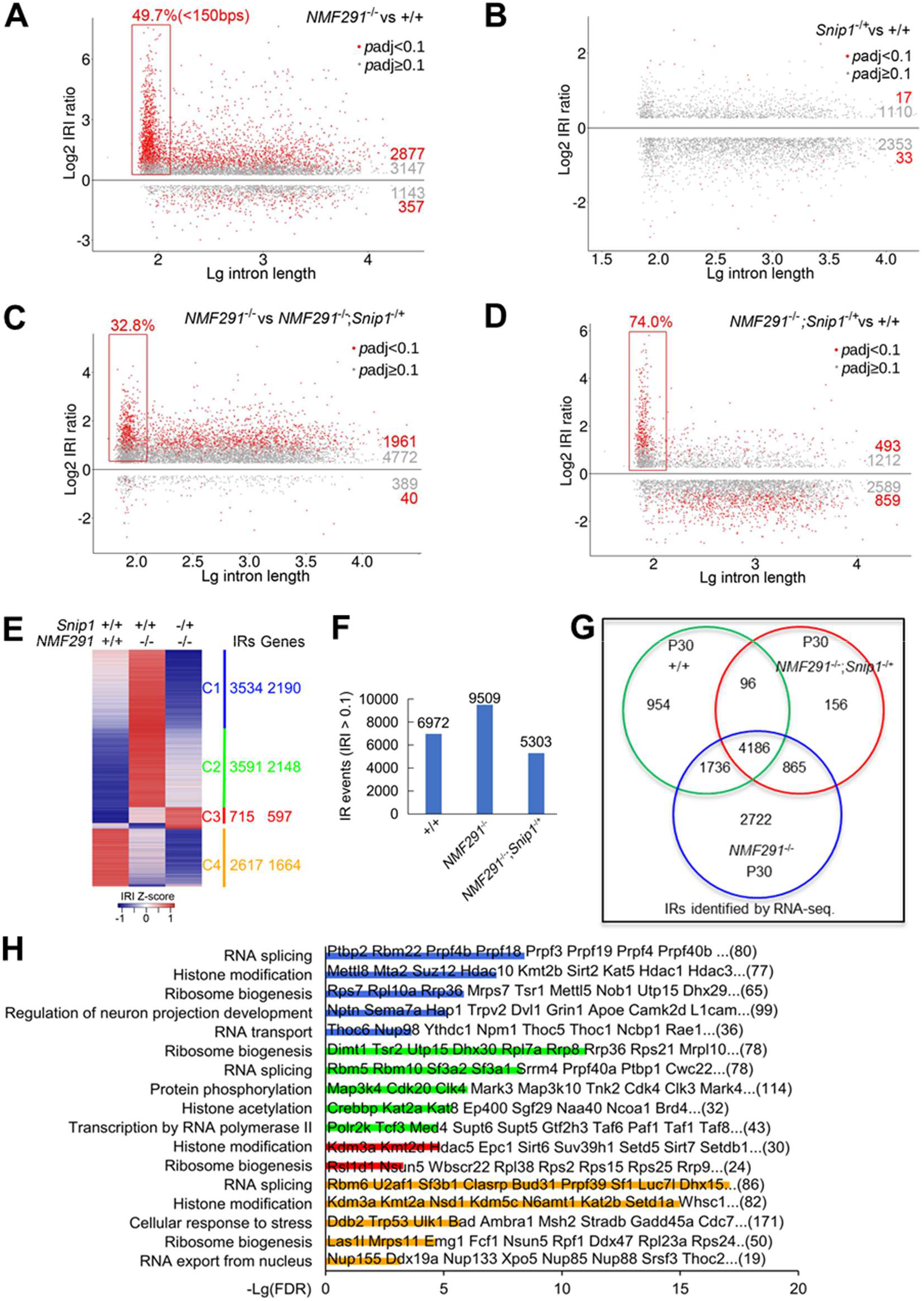
Haploinsufficiency of *Snip1* partially rescues the *NMF291* cerebellar IRs and the majority of them also exist in wildtype cerebellum. (A-D) Pairwise comparisons of cerebellar IRI between the indicated genotypes. We set two separate cutoffs colored by red and gray, respectively. Red dots: events with FDR adjusted *p*-value (*p*adj) < 0.1 and the IRI fold-change (FC) > 1.2; gray dots: *p*adj ≥ 0.1 and the IRI FC > 1.2. Only events with IRI > 0.1 were included for the comparison. The retained introns less than 150bps in length were highlighted with rectangles and their percentage of all the IRs (red rots) was labeled. Mice, male, 1 month of age, n=3. (E) Heatmap represents IRI changes across the indicated genotypes. Z-score was used to normalize IRIs in each row. The numbers of IRs and their corresponding genes (Genes) are shown. (F and G) Cerebellar IRs (IRI > 0.1, F) of the indicated genotypes and ages and their overlaps (G). (H) GO analysis of gene clusters shown in (E). The values in X axis are -Lg adjusted *p*-values. Gene numbers in each term were labeled. Some of gene names were labeled in the plot. See also Figure S2.

To gain detailed insights into these rescued IRs, we choose three representative IR events in *Nop2*, *Pias4*, and *Ptbp1*, and employed IGV (Integrative Genomics Viewer) to visualize them in wildtype (+/+), *Snip1*^-/+^, *NMF291*^-/-^, and *NMF291*^-/-^;*Snip1*^-/+^ cerebella. Indeed, *Snip1*^-/+^ largely rescued these IRs (Figs. S2C and S2D). In addition, we noticed higher expression levels of the corresponding genes in *NMF291*^-/-^ cerebella compared to that of other genotypes, suggesting that the higher gene expression was also largely rescued by *Snip1*^-/+^. To examine it globally, we compared expression level of the corresponding genes, whose IRs were rescued with high confidence (1961 events shown in Fig. 2C), in +/+, *NMF291*^-/-^, and *NMF291*^-/-^;*Snip1*^-/+^ cerebella (Fig. S2E). Gene expression levels comparing wildtype and *NMF291*^-/-^ were mutually exclusive, with upregulated genes in the *NMF291*^-/-^ cerebellum globally rescued by *Snip1*^-/+^, and little effect on downregulated genes.

To further understand the corresponding gene functions, we extracted IRs (IRI > 0.1) from the +/+ (6972), *NMF291*^-/-^ (9509), and *NMF291*^-/-^;*Snip1*^-/+^ (5303) cerebella and compared IR level changes across different genotypes (Figs. 2E and 2F). Interestingly, 62.3% and 53.1% of IRs shown in the *NMF291*^-/-^ cerebella also appear in +/+ and *NMF291*^-/-^;*Snip1*^-/+^, respectively (Fig. 2G). IR events were grouped into four clusters across the different genotypes and the corresponding genes in each cluster are related to several cellular processes (Figs. 2E and 2H). For cluster 1 (C1) genes, the IRs were fully rescued by *Snip1*^-/+^ (Fig. 3A) and the corresponding genes are highly enriched in RNA splicing, histone modification, ribosome biogenesis, neuronal projection development, and RNA transport (Fig. 2H). The IRs of cluster 2 (C2) genes were less-rescued by *Snip1*^-/+^. This group of genes are involved in ribosome biogenesis, RNA splicing, protein phosphorylation, histone modification, and transcription by RNA polymerase II. Cluster 3 (C3) genes are unique, and their IRs were not rescued but even over-represented in *NMF291*^-/-^;*Snip1*^-/+^ cerebella, although their number is less than that of other clusters. These genes are enriched in histone modification and ribosome biogenesis. The IR events in cluster 4 (C4) genes were under-represented in the *NMF291*^-/-^ cerebella compared to that of +/+, and became even less-represented in *NMF291*^-/-^;*Snip1*^-/+^. These genes are enriched in RNA splicing, histone modification, and cellular response to stress. Taken together, we demonstrate that 1) haploinsufficiency of *Snip1* globally rescues the IRs and their corresponding gene expression shown in the *NMF291* mutant cerebellum; 2) the cerebellar IRs are not randomly distributed in their transcripts; 3) the majority of IRs over- represented in the *NMF291*^-/-^ cerebella also exist in that of the wildtype; and 4) genes involved in RNA metabolism/processing and cellular response to stress tend to transcribe intron-containing transcripts.

**Figure 3.**
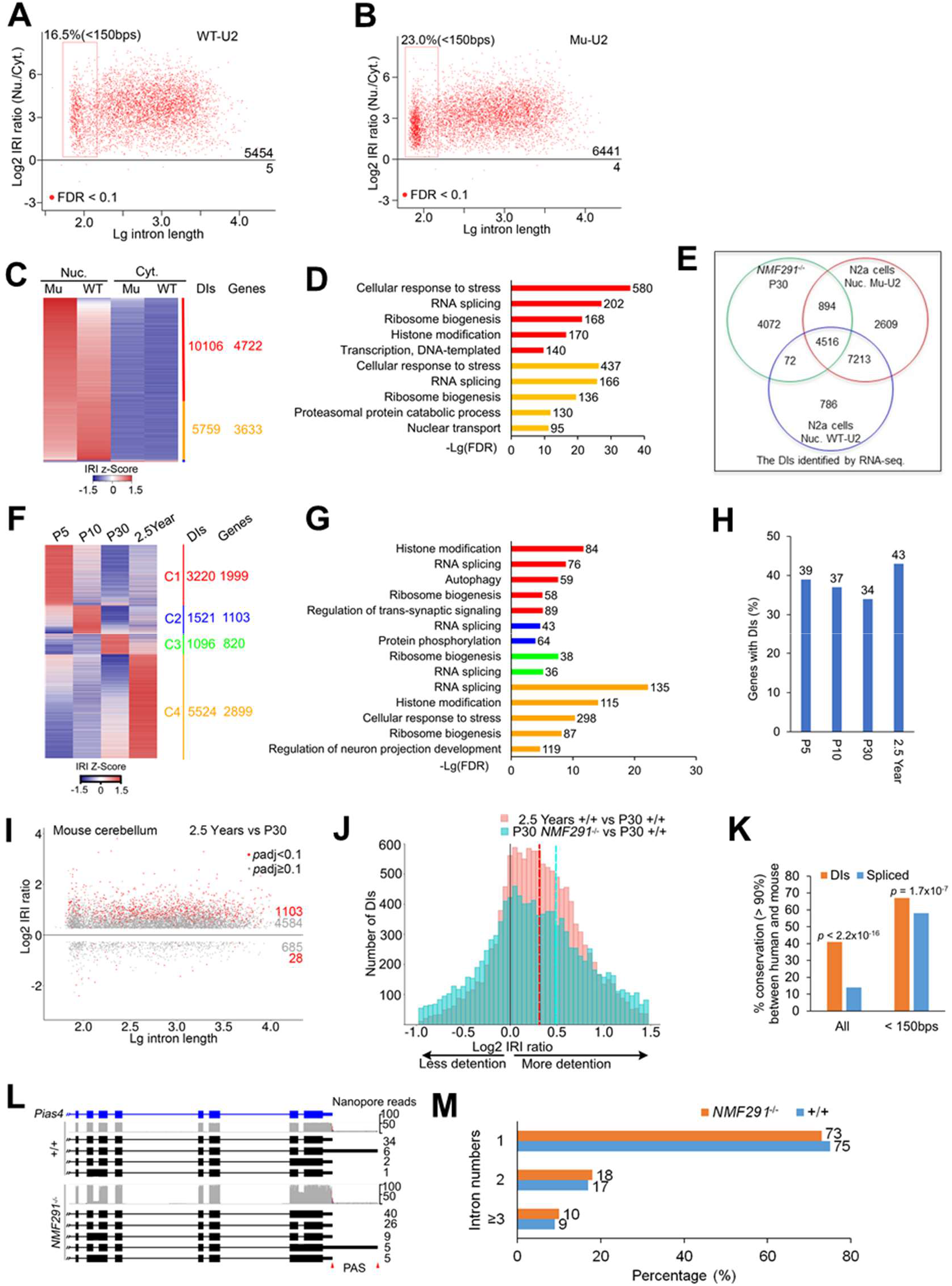
Intron-containing transcripts overrepresented in the *NMF291* mutant cerebellum are IDTs that are developmentally regulated. (A and B) The nuclear and cytosolic fractions of N2a cells expressing WT-U2 or Mu-U2 were applied for RNA-seq, respectively. The IRI ratio of nucleus to cytosol was plotted (n = 3, *p*adj < 0.1). (C and D) Heatmap represents the DIs in nucleus (Nuc.) and cytosol (Cyt.) of N2a cells expressing WT-U2 or Mu-U2. The numbers of the DIs and their corresponding genes (Genes) are shown. GO analysis (D). The X axis shows -Lg adjusted *p*-values. Gene number in each term was labeled. (E) A majority of intron-containing transcripts overrepresented in *NMF291* mutant cerebellum also appeared in nucleus extraction of N2a cells. (F and G) Heatmap represents IRI changes across different ages by RNA-seq. We included the DIs with IRI > 0.1 and clustered them. GO analysis (G). Mice, +/+, 3 males for each age point. (H) Percentage of cerebellum-expressed genes with DIs detected by RNA-seq. Wildtype P5, 10, 30, and 2.5-year mouse cerebella were applied for RNA-seq. Genes with the DI surrounding exon reads > 20 were considered as cerebellum-expressed genes. (I) Comparisons of cerebellar IRI between aged (2.5 years) and young (P30) mice. We set two separate cutoffs colored by red and gray as shown in Figure 2. Mice, male, n=3. (J) Distribution of DIs with their IRI changes in aged (2.5 years) and P30 *NMF291*^-/-^ mouse cerebellum compared to P30 wildtype control. X axis is log2 adjusted IRI ratio. Dashed lines indicate medians of log2 adjusted IRI ratio in these two groups. (K) Evolutionary conservation of the DIs (IRI > 0.1) and efficiently spliced (IRI < 0.02) introns detected in the *NMF291*^-/-^ mutant cerebellum by RNA-seq. *p* values correspond to two-sided proportion tests. (L) The representative DIs detected by nanopore-sequencing. Cerebellar mRNAs extracted from wildtype (+/+) and *NMF291*^-/-^ animals (one month of age, n=3). PAS, polyadenylation site. (M) A majority of DIs detected by nanopore-sequencing contain 1-2 intron(s) in wildtype (90%) and mutant (91%) cerebella. See also Figures S3-S5.

### Intron-containing transcripts overrepresented in mutant U2 cerebellum are IDTs featured by nuclear-localized, incompletely spliced, and polyadenylated

Intron-containing transcripts often contain PTCs, triggering NMD for degradation (*14, 31*). However, in *NMF291* mutant cerebellum, intron-containing transcripts are abundant and stable (*28*) (Figs. 2 and S2), which prompted us to examine whether these transcripts are targeted by NMD. To this end, we cultured Neuro2a (N2a) cells, a mouse neuroblastoma cell line, and transfected the cells with wildtype (WT-U2) and mutant U2 (Mu-U2) snRNA expressing plasmids. Compared to WT-U2, expression of Mu-U2 increased IRs at several endogenous sites, including *Ptbp1* intron5 (Fig. S3A). The IRs were not sensitive to inhibition of NMD, either by addition of cycloheximide (CHX) in the culture medium or knockdown of *Upf1* (Figs. S3B-S3E), suggesting that they are stable and detained in nucleus (*14, 15*). To test this possibility, we performed nuclear and cytosolic fractionation and examined IRs (Fig. S3F). Irrespective of WT- or Mu-U2 expression, IRs were enriched in the nuclear fraction. Therefore, we suggest that these incompletely spliced introns are DIs, previously featured by polyadenylated, detained in the nucleus, and immune to NMD (*15*).

To globally examine the nuclear enrichment of IDTs, we performed RNA-seq with poly(A) selection in isolated nuclear and cytosolic fractions in N2a cells (Figs. 3A and 3B). The IRI ratio of the nucleus to the cytosol indicated a global nuclear enrichment of polyadenylated IDTs in these cells regardless of whether WT-U2 or Mu-U2 was expressed. Consistent with what we observed in the *NMF291* mutant cerebellum, Mu-U2 expression in N2a cells increased the number of DIs (Figs. 3C). Genes transcribing these IDTs are functionally involved in RNA metabolism/processing and cellular response to stress (Fig. 3D), similar to what we observed *in vivo* (Fig. 2H). In fact, a majority of the DIs (56.6%) in the *NMF291* mutant cerebellum also appear in N2a cells expressing Mu-U2 (Fig. 3E).

Overrepresented IDTs in *NMF29*1 mutant cerebellum (Figs. 2 and S2) allowed us to examine whether these polyadenylated IDTs produce protein or not with high confidence. *Ptbp1* transcripts with intron 5 were a minor form in the wildtype cerebellum but became dominant in the mutant (Figs. S2C and S4A). Using a PTBP1 N-terminal antibody, we detected comparable amounts of PTBP1 (∼50kD) in wildtype and mutant cerebella but failed to detect the corresponding truncated protein supposedly produced from *Ptbp1* intron 5- containing transcripts (Figs. S4B and S4C), supporting the idea that the IDTs are detained in nucleus with limited accessibility to cytoplasmic protein translation machinery. To globally examine the protein products derived from DIs, we generated a customized peptide database, including peptides encoded by the DIs found in wildtype and *NMF291* mutant cerebellum and their upstream exons (Fig. S4D). Although we retrieved thousands of peptides coded by the upstream exons, we failed to retrieve any peptide encoded by the DIs from three biological replicates of the wildtype and *NMF291* mutant cerebellar protein lysates (Fig. S4E). Therefore, we suggest that the majority of intron-containing transcripts over-represented by expression of mutant U2 are IDTs featured by nuclear-localized, incompletely spliced, and polyadenylated.

### More than 1/3 of cerebellum-expressed genes transcribe IDTs that are highly regulated and ∼90% of these IDTs only contain 1-2 intron(s)

To test whether DIs are regulated during cerebellum development and aging, we analyzed cerebellar DIs (IRI > 0.1) at P5, 10, 30, and 2.5-year by using RNA-Seq and grouped them into four clusters (Fig. 3F). Every developmental age point has their unique IDTs transcribed by genes involved in several cellular processes, including RNA splicing, cellular response to stress, histone modification, and ribosome biogenesis (Fig. 3G). To examine how much percentage of cerebellum-expressed genes have DIs, we extracted 8917 genes with reasonable expression (the DI surrounding exon reads > 20) in P5, 10, 30, and 2.5-year cerebella (Fig. 3H). Among them, 34.2 - 43.2% have reliable DIs (IRI > 0.1) with higher percentages in P5 and 2.5-year cerebella. The IRI ratio of 2.5-year to P30 wildtype indicated an enrichment of IDTs in aged cerebellum (Fig. 3I). However, compared to the DIs overrepresented in mutant cerebellum, less detention appears in aged cerebellum, with smaller mean value of log2 IRI ratio (0.32 for aged, 0.48 for *NMF291*^-/-^ cerebellum, both compared to that of P30 +/+) (Fig. 3J). To understand the details of DIs at single transcript level, we employed nanopore sequencing, a long-read sequencing technology (*32*), to detect the DI features in P30 and 2.5-year wildtype and *NMF291*^-/-^ cerebella. To call for full length IDTs, we only included the nanopore reads containing both 5’UTR and 3’UTR with IRI > 0.05. The majority of these DIs (73%) shown in the aged cerebellum also appear in that of *NMF291* mutant cerebellum (Fig. S5A). In addition, higher percentage of IDTs of all full length transcripts we examined showed in both aged (16.0%) and *NMF291* mutant (17.5%) cerebella (Fig. S5B), compared to that of P30 wildtype. These indicate that the DIs are primarily affected during aging process, presumably when the function of spliceosome declines.

Next, we asked whether these DIs are evolutionarily conserved between mouse and human. We categorized the efficiently-spliced introns (IRIs < 0.02) and the DIs (IRIs > 0.1) in the *NMF291*^-/-^ mutant cerebella. Compared to the efficiently-spliced introns, the DIs are significantly more conserved between human and mouse (p < 2.2 x 10^-16^) (Fig. 3K). For intron length of less than 150 bps, DIs also showed significantly more conserved (p < 1.7 x 10^-7^) than that of efficiently-spliced introns.

To further learn the DI features at full length transcript level, we retrieved 177002 IDTs for wildtype (+/+) and 334999 for *NMF291* mutant (*NMF291*^-/-^) cerebella by using nanopore sequencing. Consistent with our 2^nd^ generation RNA-seq results, nanopore-reads showed more IDTs in mutant cerebellum than that of wildtype (Fig. 3L). However, when we categorized the IDTs in terms of their intron number distribution, most IDTs contained 1-2 intron(s) in wildtype (91%) as well as in mutant (90%) cerebella (Fig. 3M). In addition, the percentage of intron number distribution between the two genotypes is almost identical with ∼70% IDTs containing only one intron in both genotypes. Taken together, our findings reveal that 1) over 1/3 of cerebellum-expressed genes transcribe IDTs and their DI splicing is highly regulated during cerebellum development and aging; 2) evolutionarily these DIs are more conserved than that of efficiently-spliced; and 3) ∼90% of IDTs only contain 1-2 intron(s), indicating that DIs only comprise a small proportion of the total introns transcribed in cerebellum.

### Nuclear-localized SNIP1 binds to cerebellar polyadenylated IDTs

Previous studies showed that SNIP1 interacts with the TGF-β family Smad proteins, NF-κB transcription factor p65, and transcriptional coactivators p300/CBP, resulting in regulation of TGF-β and NF-κB signaling (*33, 34*). Post-transcriptionally, SNIP1 also regulates Cyclin D1 RNA stability (*35*). However, how SNIP1 regulates pre-mRNA splicing *per se* are not studied in mammal. If DI splicing is regulated by SNIP1, we speculated that SNIP1 must bind to these transcripts. To this end, we inserted 3 × Flag tag at the C-terminal end of *Snip1* by Crispr-Cas9-mediated homologous recombination (Figs. 4A, S6A, and S6B). SNIP1-Flag appeared in the knockin (KI) cerebellum at the expected molecular weight, and was absent in the wildtype control (Fig. 4A). The cerebellar expression level of SNIP1-Flag at P5 and P10 was 5.0 and 3.6 times higher than that of P30 (Fig. 4A). Ubiquitous expression of SNIP1 was documented in various adult mouse tissues (Fig. S6C).

**Figure 4.**
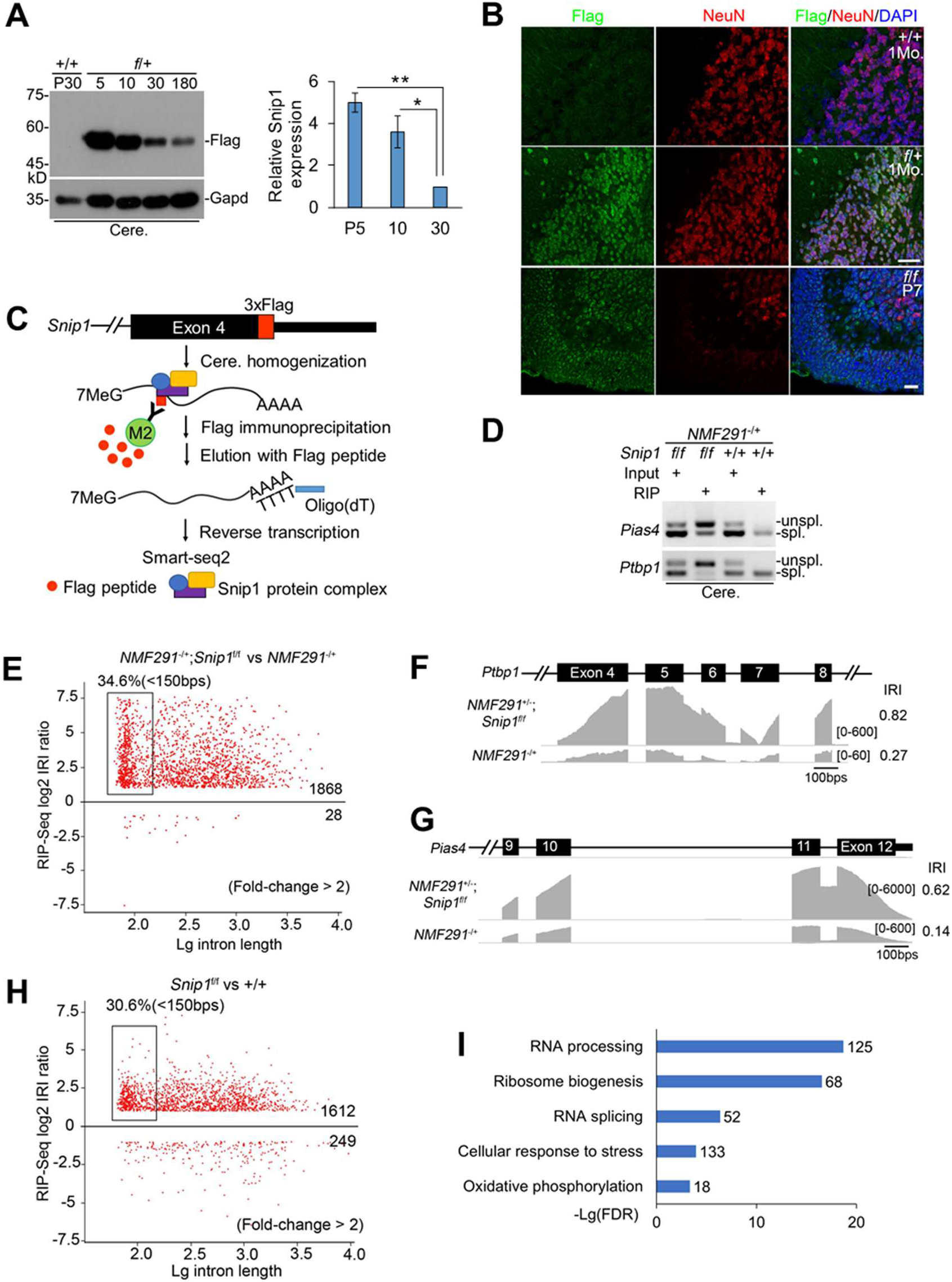
Nucleus-localized SNIP1 binds to cerebellar IDTs. (A) The expression level of SNIP1 was documented with a Flag antibody at various age points in *Snip1-Flag* KI animals. GAPDH, loading control; *f*/+, mice heterozygous for *Snip1-Flag* KI; +/+, negative control for Flag immunoblot. A summary of the expression level was inserted. (B) Flag-immunostaining was performed in *Snip1-Flag* KI mouse cerebellum at one month and P7 of ages. NeuN, a neuronal marker; +/+, negative control for Flag immunostaining. Scale bar for one month, 30 μm; for P7, 20 μm. (C) A diagram for native Flag-RIP-seq. The RNA/protein complex containing Flag-SNIP1 was eluted by competition with free Flag peptides. The resulting Flag-SNIP1-bound RNAs were applied for Smart-seq2 to amplify full-length polyadenylated mRNAs. M2, M2-beads conjugated with Flag-antibody. (D and E) RIP was performed with cerebellar protein lysates from *NMF291*^-/+^;*f*/*f* or *NMF291*^-/+^ mice at one month of age. Enrichment of DIs was detected by RT-PCR in RIP products from *NMF291*^-/+^;*f*/*f* mice but not from that of *NMF291*^-/+^ controls. Global examination of the enrichment by Flag-RIP-seq from three biological replicates (E). (F-G). Two representative SNIP1-bound IDTs visualized by IGV. (H and I) Global examination of enrichment of SNIP1-bound DIs in P7 *Snip1^f^*^/*f*^ cerebellum by Flag-RIP-seq (n = 3). Wildtype (+/+) animals served as negative control for Flag-RIP-seq. GO analysis of the genes transcribing SNIP1-bound IDTs shown in (I). In A, the values are presented as mean ± SEM (mice, n ≥ 3 for each age point). \**p*<0.05, \*\**p*<0.01, ANOVA, SPSS. See also Figure S6.

Nuclear Flag-immunoreactive signals appeared in *Snip1-Flag* KI adult and P7 cerebella, and were absent in wildtype control (Fig. S6D). In the adult cerebellum, the majority of SNIP1-Flag signals were NeuN-positive in the internal granule layer (IGL), indicating that SNIP1 is majorly expressed in adult cerebellar granule cells. In P7 cerebellum, the Flag-immunoreactive signals were evident in proliferative granule cell progenitors in the external granule layer (EGL), migrating granule cells in the molecular layer (ML), and NeuN-positive granule cells in the IGL (Fig. 4B).

To examine whether SNIP1 binds to the post-transcriptional polyadenylated IDTs in *NMF291* mutant cerebellum, we employed a native RNA immunoprecipitation (RIP) coupled with poly(A) selection strategy (Fig. 4C). Instead of crosslinking RIP, native RIP enables us to detect entire transcripts in a more stable and long-lasting RNA-protein complex. Native Flag-RIP precipitated *Pias4* and *Ptbp1* IDTs in *NMF291* mutant cerebella expressing SNIP1-Flag (*NMF291*^-/+^;*Snip1^f^*^/^*^f^*), which were depleted in that of *NMF291*^-/+^ without expression of SNIP1-Flag (Fig. 4D). To globally examine the binding, we performed native RIP followed by Smart-seq2 (*36*), which amplifies full-length polyadenylated mRNAs (Fig. 4C). The IRI ratio of *NMF291*^-/+^;*Snip1^f^*^/^*^f^* to *NMF291*^-/+^ revealed an enrichment (1868 vs 28) of DIs (Fig. 4E), which was also visualized in two representative *Ptbp1* and *Pias4* IDTs by IGV (Figs. 4F and 4G). Among these transcripts, 34.6% of them were small introns with a size of less than 150bps (Fig. 4E).

To examine whether the binding is independent of mutant U2 expression, we performed Flag-RIP-seq in *Snip1-Flag* KI animals at P7, when SNIP1 is highly expressed (Fig. 4H). The IRI ratio of *Snip1^f^*^/^*^f^* to wildtype (1612 vs 249) suggested the binding of SNIP1 to the IDTs in P7 cerebella (Fig. 4H). Among them, 30.6% are small introns with a length of less than 150 bps. Genes transcribing these transcripts are involved in RNA processing, ribosome biogenesis, RNA splicing, oxidative phosphorylation, and DNA repair (Fig. 4I). Therefore, we conclude that nuclear-localized SNIP1 deposits at a group of cerebellar IDTs encoded by genes involved in several cellular processes, especially RNA metabolism/processing and cellular response to stress.

### DI splicing is paused prior to the first catalytic step

To understand how SNIP1 regulates DI splicing, we examined SNIP1 protein binding partners in N2a cells, a cellular model that carries endogenous DIs (Fig. S3). In N2a cells stably expressing SNIP1-Flag, we performed Flag-co-immunoprecipitation (co-IP) followed by mass spectrometry (co-IP/MS). High confidence hits from three biological replicates were protein components found in spliceosome and peripheral EJC, or proteins involved in RNA export (Fig. 5A and Table S3). These SNIP1-interacting protein partners include RNA helicases (BRR2 and SNU114) and SR/SR-like proteins (SFRS16, SRM300, ACIN1, RNPS1, and SRSF7). We noticed that protein components found in spliceosome are that of B^act^, the activated spliceosome, but not that of B*, the catalytically activated spliceosome (*7, 37, 38*) (Fig. 5B). Given that SNIP1 binds to IDTs (Fig. 4) and recent resolved human spliceosome structure also support the presence of SNIP1 in B^act^ spliceosome but not B and C complex (*7, 37, 39–41*), we proposed that DI splicing is paused at B^act^.

**Figure 5.**
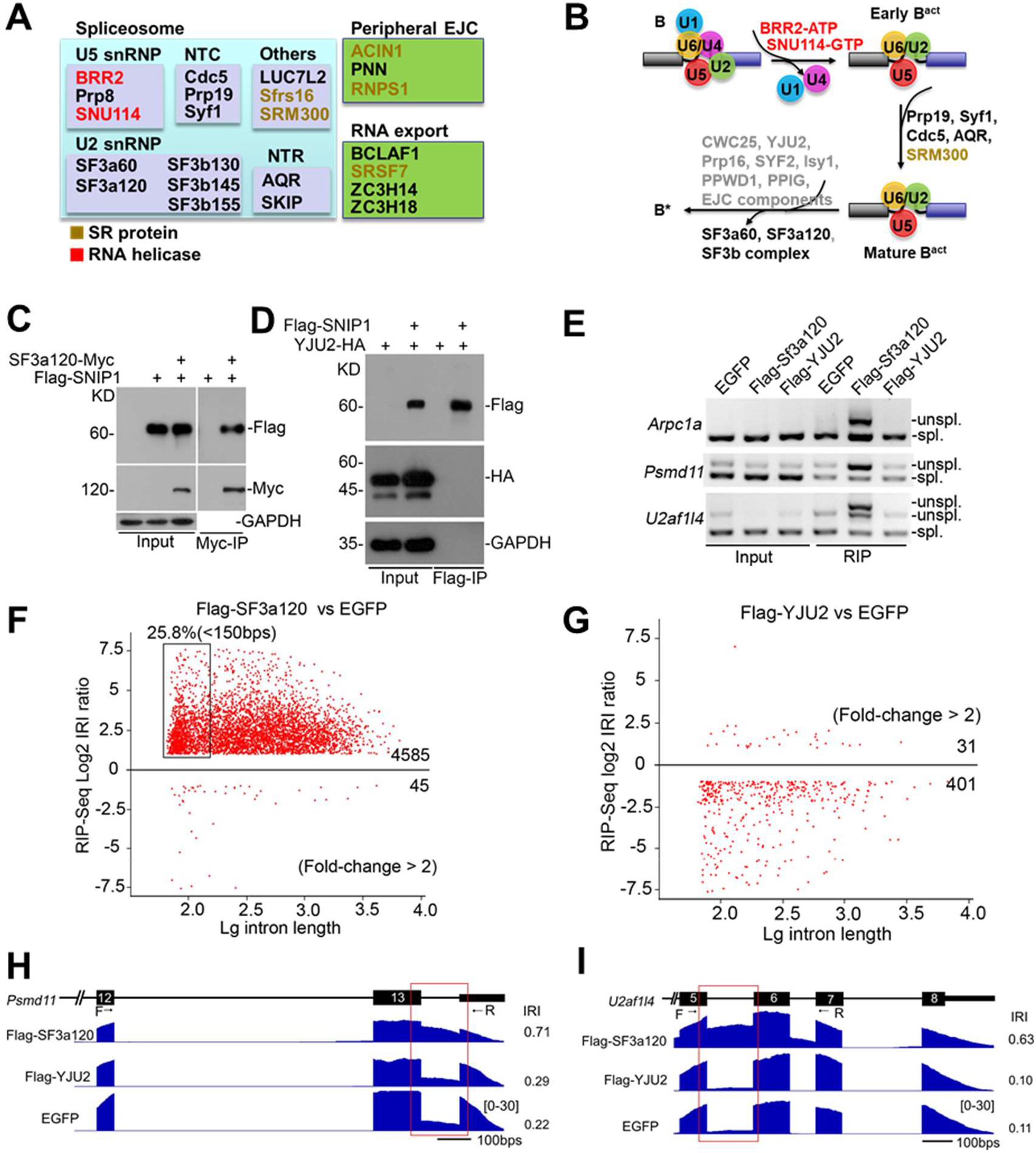
DI splicing is likely paused at B^act^. (A) Potential SNIP1 interacting proteins identified by Flag-IP coupled with mass spectrometry (co-IP/MS) in N2a cells expressing Flag-SNIP1. Many of these protein partners are involved in pre-mRNA splicing and RNA metabolism (also see Table S3). (B) The potential SNIP1 interacting partners are found in B^act^ but not B*. Factors loaded into B* are labeled in gray, which were not hit by our co-IP/MS. (C and D) Interactions between SNIP1 and protein components found in B^act^ component SF3a120 (C) but not those in B* component YJU2 (D) were validated in N2a cells expressing the indicated tagged proteins. (E) RIP was performed in N2a cells expressing Flag-SF3a120, Flag-YJU2, or EGFP. DIs were measured by RT-PCR using Smart-amplified cDNA. (F-I) Flag-RIP-seq was performed in N2a cells expressing Flag-SF3a120 (F) and Flag-YJU2 (G). N2a cells expressing EGFP, a negative control for our Flag-RIP-seq. Two representative SF3a120-bound DIs visualized by IGV (H and I). See also Table S3, Figure S7.

Release of SF3a and SF3b complexes is a key step for B^act^ to B* transition (*5*). Interactions between SNIP1 and SF3a or SF3b components, including SF3a60, SF3a120, SF3a66, and SF3b130, were validated in N2a cells expressing tagged SNIP1 (Figs. 5C and S7A-S7C). Although interaction between SNIP1 and Syf1, another B^act^ component, was further confirmed in the N2a cells, we failed to detect interactions between SNIP1 and B* components, including YJU2 and CWC25, in the similar conditions to those for B^act^ components (Figs. 5D and S7D-S7E). As EJC core proteins, including eIF4AIII, MAGOH, Y14, and MLN51, are recruited to the B* and C complex (*8, 41, 42*), we failed to detect the interaction between SNIP1 and eIF4AIII (Fig. S7F).

To examine whether B^act^ is indeed found at DIs, we expressed Flag-tagged SF3a120, a B^act^ component, and YJU2, a B* and C component and step-I specific factor (*7, 37–39, 41*), in N2a cells and performed Flag-RIP. SF3a120 precipitated the IDTs, like *Arpc1a*, *Psmd11*, and *U2af1l4*, which were not enriched by YJU2 or EGFP controls (Fig. 5E). To globally test the association of SF3a120 with IDTs, we performed Flag-RIP-seq in N2a cells expressing Flag-SF3a120, Flag-YJU2, or EGFP. The IRI ratio of Flag-SF3a120 to EGFP showed an enrichment (4585 vs 45) of IDTs (Fig. 5F). In contrast, the IRI ratio of Flag-YJU2 to EGFP indicated no such enrichment (31 vs 401) (Fig. 5G). The enrichment was visualized by IGV in two representative DI events (Figs. 5H and 5I). Therefore, we suggest that DI splicing is likely paused at B^act^ prior to the first catalytic step of pre-mRNA splicing.

### SNIP1 and RNPS1 function as molecular brake for spliceosome pausing at DIs

As we documented above, haploinsufficiency of *Snip1* partially rescues DIs from accumulating in *NMF291* mutant cerebellum (Fig. 2). In N2a cells, knockdown of *Snip1* by shRNA (sh-Snip1) also reduced the levels of several endogenous intron detention events that were amplified by the expression of Mu-U2 (Figs. S8A and S8B). In addition to protein components found in B^act^, protein components of the peripheral EJC (ACIN1, PNN, and RNPS1) were also identified as potential SNIP1 interacting partners (Table S3). Like *Snip1*, knockdown of *Pnn* and *Rnps1* significantly reduced the intron detention amplified by Mu-U2 (Figs. 6A and 6B and S8C). Interaction of SNIP1, PNN, and RNPS1 was confirmed by co-IP in N2a cells simultaneously expressing tagged PNN, SNIP1, and RNPS1 (Fig. S8D), suggesting that these proteins work together to regulate spliceosome pausing. Although splicing factor SRm300 was found by our co-IP/MS (Table S3), SRm300 knockdown did not significantly influence the intron detention we examined (Figs. 6A and 6B).

**Figure 6.**
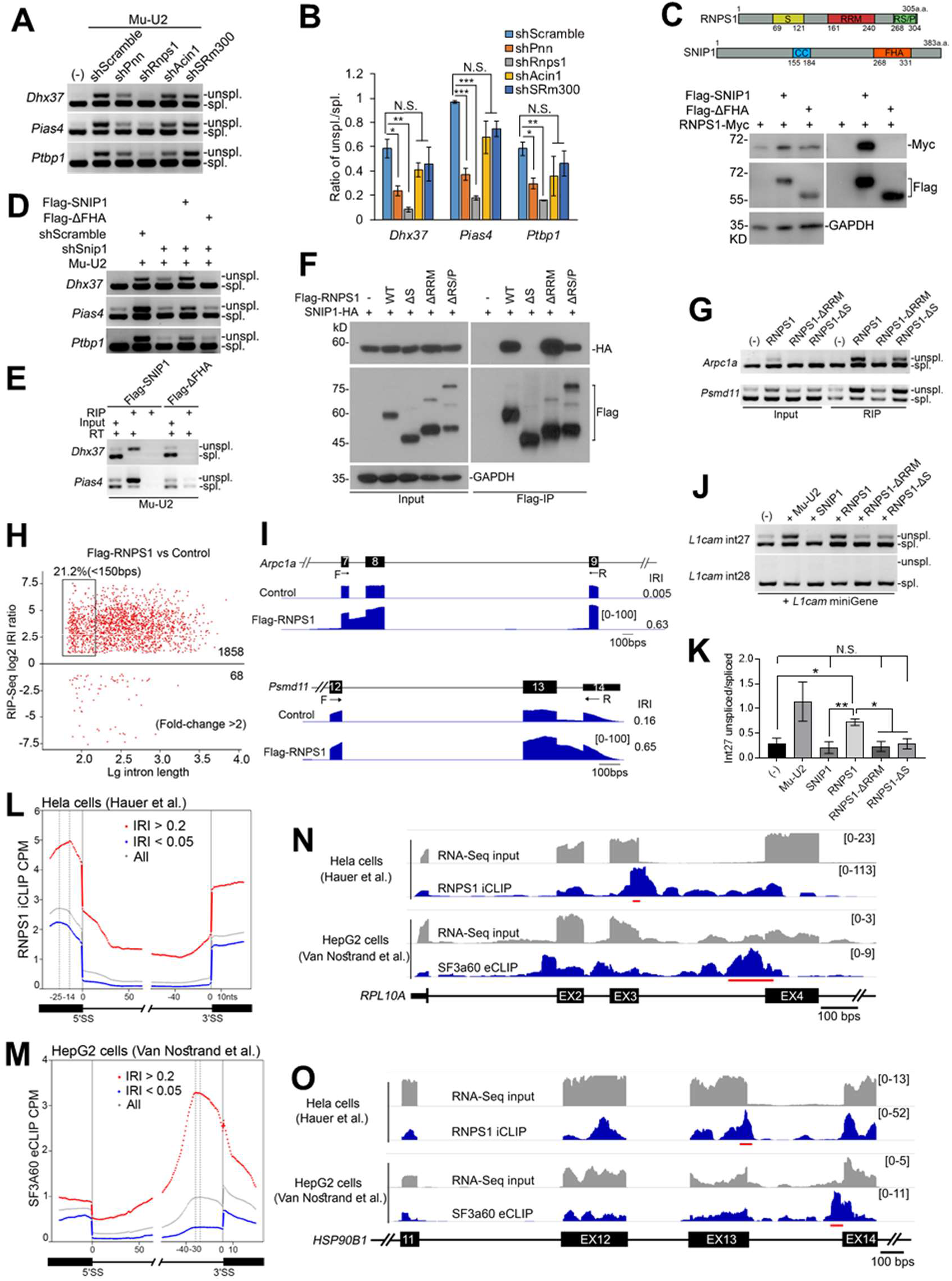
SNIP1 and RNPS1 function as a molecular brake for spliceosome pausing at DIs. (A and B) Effect of the knockdowns of genes encoding peripheral EJC components (*Acin1*, *Pnn*, and *Rnps1*) and a splicing factor (*SRm300*) on the DIs (A) amplified by expression of mutant U2 (Mu-U2) in N2a cells and the data summary (B). (C) Annotated domains of SNIP1 and RNPS1 (upper). Co-IP was performed in N2a cells expressing the Flag-tagged full-length or FHA domain truncated SNIP1 (ΔFHA) together with RNPS1-Myc (lower). (D) N2a cells were infected with scrambled shRNA or shSnip1 and transfected with the indicated expression plasmids. (E) Flag-RIP was performed with protein lysates from N2a cells infected with Flag-tagged full-length or ΔFHA SNIP1 and transfected with Mu-U2 expression plasmid. RT, reverse transcriptase. (F) Co-IP was performed in HEK293 cells expressing HA-SNIP1 and/or Flag- tagged RNPS1, including full length and various domain-truncated RNPS1. (G) Flag-RIP was performed with protein lysates from N2a cells expressing the indicated proteins. Smart-amplified cDNA was analyzed by RT-PCR. (H and I) Global examination of the RNPS1-bound DIs in N2a cells expressing Flag-RNPS1. Control, naïve N2a cells. Two representative RNPS1-bound DIs visualized by IGV (I). (J and K) HEK293 cells were transfected with previously described splicing minigene reporters derived from *L1cam* detained intron 27 (int27) and constitutively spliced intron 28 (int28) (PMID: 22265417). The data summary was shown in (K). (L and M) CLIP-seq read density plot for RNPS1 (L) and SF3a60 (M). Read density plot with DIs (IRI > 0.2) shown in red; read density plot with efficiently- spliced introns (IRI < 0.05) shown in blue; read density plot regardless of IRI shown in gray. RNPS1 iCLIP data from ArrayExpress (PMID: 27475226) and SF3a60 eCLIP data from ENCODE (PMID: 27018577). (N and O) Read density tracks along two representative genes *RPL10A* (N) and *HSP90B1* (O) viewed by IGV and scaled by RPM (reads per million usable). The most significant clusters called by CLIPper and illustrated by the red lines. In A, D, E, G, and J, DIs were measured by RT-PCR. Unspl., unspliced transcripts; spl., spliced transcripts. In B and K, the values are presented as mean ± SEM, n=3. \**p*<0.05, \*\**p*<0.01, \*\*\**p*<0.001, N.S., no statistical significance, ANOVA, SPSS. See also Figure S8.

Interaction between SNIP1 and RNPS1 depends on SNIP1 FHA (forkhead-associated) domain, a small protein module involved in phospho-dependent protein/protein interaction (*43, 44*), because deletion of the FHA domain (Flag- ΔFHA) abolished the SNIP1-RNPS1 interaction (Fig. 6C). Exogenously expressed full-length SNIP1 but not ΔFHA SNIP1 restored intron detention amplified by Mu-U2 in N2a cells infected with shSnip1, suggesting that interaction between SNIP1 and RNPS1 is required for spliceosome pausing at DIs (Figs. 6D and S8E). In addition, unlike Flag-SNIP1, Flag-ΔFHA failed to precipitate IDTs (Fig. 6E), suggesting that SNIP1 binds to the IDTs through interaction with RNPS1.

To examine how RNPS1 interacts with SNIP1, we expressed full-length and various truncated forms of RNPS1, including ΔS (a serine-rich domain deletion), ΔRRM (an RNA recognition motif deletion), and ΔRS/P (an arginine and serine/proline-rich domain deletion), together with SNIP1 in N2a cells (Fig. 6F). ΔS abolished RNPS1-SNIP1 interaction but ΔRRM and ΔRS/P did not, suggesting that RNPS1 interacts with SNIP1 through its serine-rich domain.

Unlike RNPS1, SNIP1 does not contain an annotated RNA binding domain. The RRM of RNPS1 is involved in formation of peripheral EJC complexes, which in turn facilitate RNA binding (*45, 46*). Flag-RNPS1 precipitated IDTs in N2a cells expressing Mu-U2, and this was absent in control cells not expressing the Flag-RNPS1 (Fig. S8F). In addition, RNPS1 binds to IDTs independent of Mu-U2 expression (Fig. 6G). However, the binding was abolished by RNPS1- ΔRRM but not RNPS1-ΔS, suggesting that RRM but not interaction between SNIP1 and RNPS1 is required for the binding. Our Flag-RIP-seq data further support the idea that RNPS1 binds to IDTs (Figs. 6H and 6I). Among the RNPS1-bound DIs, 21.2% of them are small introns with a length of less than 150bps (Fig. 6H).

If interaction between SNIP1 and RNPS1 is required for spliceosome pausing at highly regulated DIs, we assumed that ΔS RNPS1 would lose its ability to modulate the pausing. To this end, we employed two previously reported splicing reporters derived from two neighboring *L1cam* introns, one detained intron 27 (int27) and one constitutively spliced intron 28 (int28) (*28*). Severe intron detention of int27 shown in the *NMF291* mutant cerebellum was rescued by *Snip1*^-/+^ (Fig. S2B), suggesting that SNIP1 and RNPS1-containing complex modulates spliceosome pausing at int27. As previously reported (*28*), expression of Mu-U2 decreased the splicing efficiency of int27 but did not affect that of int28 constitutive splicing (Figs. 6J and 6K). Expression of full-length RNPS1 but not SNIP1 significantly decreased int27 splicing but did not affect int28 splicing, suggesting that RNPS1 is sufficient to induce intron detention. However, both ΔS and ΔRRM abolished RNPS1-mediated intron detention. Therefore, we suggest that RNPS1 docks at DIs through its RRM to induce RNPS1-SNIP1 interaction, which in turn functions as molecular brake to pause spliceosome at highly regulated DIs.

### RNPS1 and SF3a60 dock at different positions of DIs

EJC core proteins and peripheral EJC component RNPS1 have been documented to bind to mRNA and involved in post-transcriptional mRNA processing (*46–51*). To examine the binding sites of RNPS1 and SF3a complex at DIs, we reanalyzed previous reported RNPS1- and SF3a60- cross-linking and immunoprecipitation (CLIP) data (*52–54*). Globally, RNPS1 CLIP-seq reads piled up at 5’-exons (with a binding peak at -14 nts of 5’SS) and 5’-introns (between 0 to 30 nts of 5’SS) of the DIs (IRI > 0.2) (Fig. 6L), slightly differing from the binding site of core EJC (−24 to −20 nts from the 5’SS) (*47, 54*). For SF3a60 - a protein component of the SF3a complex - the CLIP-seq reads concentrated between -40 to 0 nts of the 3’SS of DIs, with a peak at -30 nts (Fig. 6M). However, such RNPS1- and SF3a60-CLIP-seq peaks were less represented at the positions of the efficiently spliced introns (IRI < 0.05) or all the introns we examined (All).

For representative CLIP peaks at DIs, IGV illustrated that intron 4 of *RPL10A* and intron 13 of *HSP90B1* are the highest confidence DIs compared to neighboring introns in both Hela and HepG2 cells (Figs. 6N and 6O). The most stringent RNPS1 binding peaks identified by CLIPper appeared at the 5’ exon/intron boundary of the highest confidence DIs. The SF3a60 peaks were located inside of the highest confidence DIs close to the 3’SS. Taken together, our analysis suggests that RNPS1 a peripheral EJC component and SF3a60 an SF3a complex component preferentially dock at DIs.

### Conditional KO of *Snip1* in cerebellar granule cells leads to IDT accumulation and neurodegeneration

SNIP1 is expressed in granule cells in the developing and adult cerebellum (Figs. 4A and 4B) and homozygous *Snip1* KO leads to embryonic lethality (Table S2). To reveal the biological consequences of *Snip1* KO in the cerebellum, we generated a *Snip1* floxed mouse line and crossed it to *Nse-CreER^T2^*, a tamoxifen-inducible Cre line specific for Cre expression mainly in cerebellar granule cells and a few granule neurons in the hippocampal dentate gyrus (Figs. S9A and S9B) (*55*). We confirmed the specificity of this system by crossing the *Nse-CreER^T2^* to a Cre-dependent reporter line, *H2B mCherry* (*56*) (Fig. S9C). After tamoxifen injections on P3, P4 and P5, we observed a deletion of *Snip1* exon 3 at both the genomic DNA and mRNA levels, specifically in the cerebellum and hippocampus but not in the cortex of *Snip1^fl^*^/^*^fl^*;*NseCreER^T2^* mice on P9 (Fig. S9D). Such deletions did not appear in the brain regions in age-matched *Snip1^fl^*^/^*^fl^* controls. We harvested P7 cerebella from *Snip1^fl^*^/^*^fl^*;*NseCreER^T2^* and age-matched *Snip1^fl^*^/^*^fl^* mice after tamoxifen injections and performed RNA-seq (Fig. 7A). Accumulation of intron-containing transcripts were evident in the *Snip1* conditional KO (cKO) cerebella compared to controls. Genes transcribing these transcripts are functionally involved in histone modification, DNA repair, RNA splicing, ribosome biogenesis, and synaptic vesicle transport (Fig. 7B), similar to what we observed in the *NMF29*1 mutant cerebellum (Fig. 2H). To understand the details of incompletely-spliced introns induced by *Snip1* cKO at the single transcript level, we sent the cKO cerebella for nanopore sequencing. Similar to what we observed in wildtype and *NMF291* mutant cerebella (Figs. 3K-3L), most intron-containing transcripts had one or two intron(s) in both *Snip1* cKO (for one, 74%; for two, 16%) and an age-matched control (*Snip1^fl^*^/^*^fl^*, for one, 78%; for two, 17%) (Fig. 7C). These data suggest that like the *NMF291* mutation, *Snip1* cKO in cerebellar granule cells leads to accumulation of IDTs, but has less effect on constitutive splicing. Indeed, the majority of intron-containing transcripts (65.7%, 3959/6026) present in *Snip1* cKO cerebella on P7 are also found in *NMF291* cerebella on P30 (Fig. 7D).

**Figure 7.**
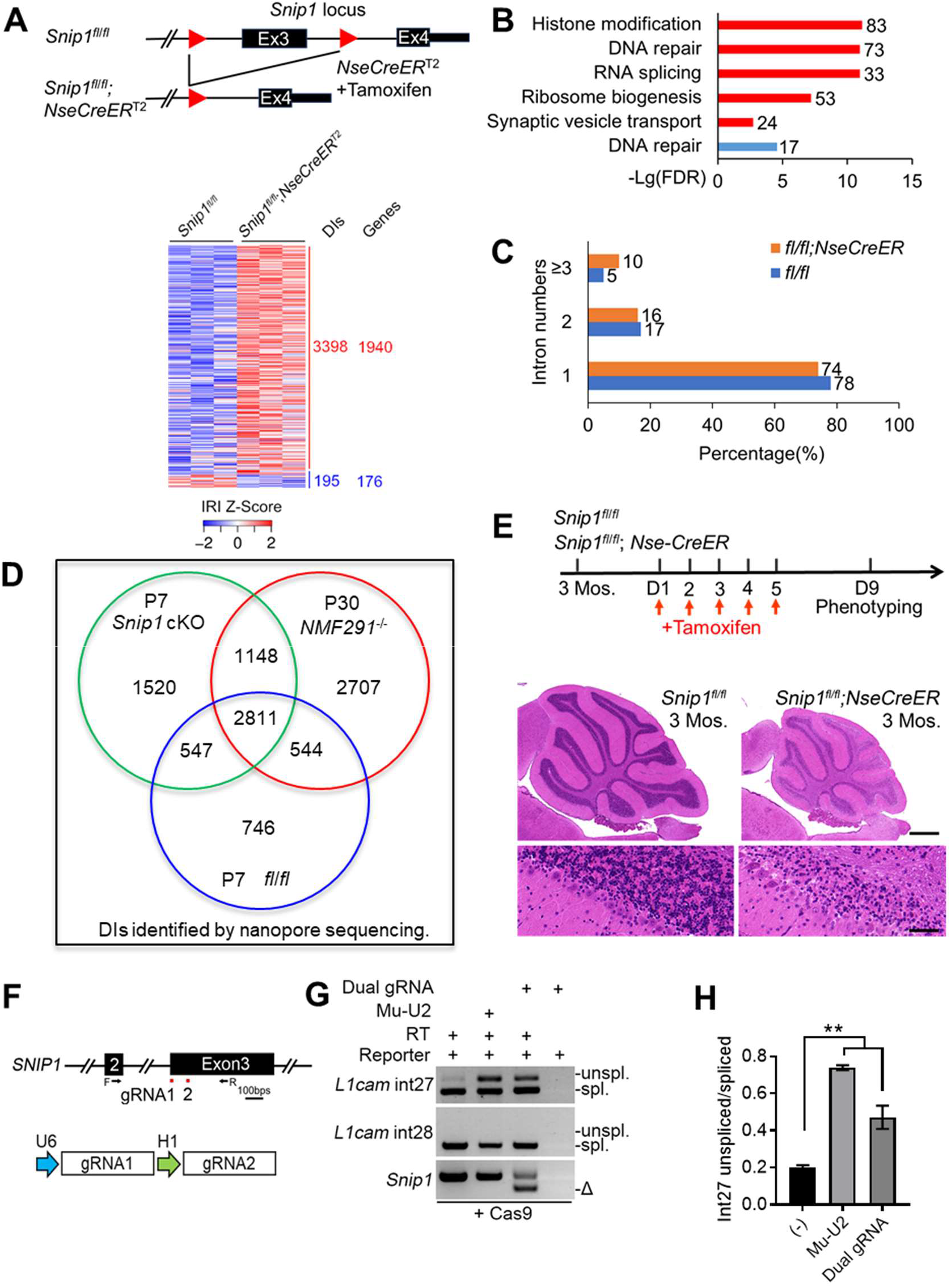
*Snip1* cKO in cerebellar granule cells leads to IDT accumulation and neurodegeneration. (A) A schematic diagram for Cre-dependent *Snip1* cKO (upper). Cerebellar RNAs extracted from the indicated genotypes on P7 (n = 3) were applied for RNA-seq. Events with IRI > 0.1 were included for clustering. (B) GO analysis of genes transcribing intron-containing transcripts that are over-represented in *Snip1* cKO cerebellum. (C) The cerebellar RNAs described in (A) were applied for nanopore sequencing. Intron number distribution of the indicated genotypes was plotted. (D) Intron-containing transcripts detected by nanopore sequencing in P7 *Snip1* cKO cerebellum are shared by that in P30 *NMF291*^-/-^ cerebellum. (E) Our experimental procedure: tamoxifen injections on day 1 (D1), 2, 3, 4, and 5 after 3 months (Mos.) of age; mouse phenotyping on D9 (upper). Hematoxylin and eosin-stained cerebellar sagittal sections of the indicated genotypes (lower). (F) Previously reported DECKO (Double Excision CRISPR Knockout, PMID: 26493208) approach was adopted for *Snip1* KO in N2a cells. *Snip1* dual gRNAs target the exon 3 (upper). The dual gRNAs are driven by U6 and H1 promoters, separately (lower). F and R, primers for Crispr/Cas9-mediated editing site fusion. (G and H) N2a cells expressing *L1cam* minigenes were transfected with Mu- U2 or *Snip1* dual gRNAs plasmids. Cas9 was constitutively expressed in these cells. DIs were measured by RT-PCR. Unspl., unspliced transcripts; spl., spliced transcripts. Δ, Crispr/Cas9-mediated editing site fusion at RNA level measured by RT-PCR. Primers used for fusion detection were labeled in (F). The data summary shown in (H). In E, the corresponding high magnification images from lobule II. Scale bar, upper 200 μm; lower, 100 μm. In H, data are presented as mean ± SEM, \*\**p*<0.01, n = 4, one-way ANOVA, SPSS. See also Figure S9 and S10.

As IDT accumulation and neurodegeneration evidenced in the *NMF291* mutant cerebellum (*28*) and the IDTs largely overlapped between *Snip1* cKO and *NMF291*^-/-^ cerebella (Fig. 7D), we then asked whether *Snip1* cKO in adult cerebellum also leads to neurodegeneration. To this end, we injected tamoxifen into adult *Snip1^fl^*^/^*^fl^*;*NseCreER^T2^* animals and age-matched controls (Fig. 7E). Nine days after injection we observed massive granule cell loss in all cerebellar lobules. To examine whether *Snip1* KO like the *NMF291* mutation impairs splicing efficiency of DI, we employed the *L1cam* splicing reporters (*28*). In order to achieve *Snip1* KO in N2a cells, we employed dual-gRNA approach (*57*) and the high fusion rate of two gRNA cutting sites at RNA level was evidenced by RT-PCR (Figs. 7F and 7G). Both Mu-U2 expression and *Snip1* KO in N2a cells significantly reduced splicing efficiency of int27 but had little effect on that of constitutive splicing of int28 (Figs. 7G and 7H). Taken together, we demonstrate that *Snip1* cKO in cerebellar granule cells cell-autonomously leads to neurodegeneration and *Snip1* KO impairs splicing efficiency at highly-regulated DIs but not constitutively-spliced introns.

## Discussion

Here, we describe interaction between SNIP1 and RNPS1 as a molecular brake for spliceosome pausing at highly regulated DIs. Together with a previous report (*28*), we suggest that misregulation of this process contributes to pathogenesis of neurodegeneration.

The yeast homolog of SNIP1 is Pml1 (pre-mRNA leakage protein 1). Compared to human SNIP1 (396 a.a.), yeast Pml1 (204 a.a.) lacks 1-180 a.a. N-terminus and has as low as ∼30% similarity to human SNIP1. Yeast RES complex contains Snu17, Bud13, and Pml1, corresponding to human RBMX2, BUD13, and SNIP1 (*58*). RES complex is initially recruited to the B/B^act^ complex and released from B* (*6, 7, 38, 44, 59-61*). We demonstrate that SNIP1 interacts with protein components found in B^act^ but not B* (Figs. 5A-5D). In yeast, unlike spliceosome basal components, such as SF3b subunits, the RES complex is not essential for yeast growth (*58, 62*). In zebrafish, individual KO of *bud13*, *snip1*, and *rbmx2* leads to mis-splicing a subset of introns (*63*). In mammalian cells, BUD13 binds to a group of retained-introns and regulates their splicing (*64*). It is plausible that SNIP1 functions together with the other two RES complex proteins to regulate spliceosome pausing at DIs, although our SNIP1 co-IP/MS experiments did not achieve reliable RBMX2 and BUD13 hits (Table S3). Given that histone modification plays an important role in alternative splicing regulation and SNIP1 regulates p300 histone acetyltransferase activity (*33, 34, 65, 66*), SNIP1 could also indirectly regulate DI splicing.

Previous studies demonstrated that RNPS1 interacts with NMD machinery for mRNA surveillance (*54, 67–69*). RNPS1 interacts with SAP18, ACIN1 or PNN to form the heterotrimeric apoptosis and splicing-associated protein (ASAP) complex (ACIN1, SAP18 and RNPS1) or the PSAP complex (PNN, SAP18 and RNPS1) (*45, 70, 71*). Both ACIN1 and PNN contain a similar RNPS1-SAP18-binding (RSB) motif to interact with RNPS1 and SAP18, suggesting that the formation of the ASAP and PSAP complexes is mutually exclusive and the resulting complexes determine the specificity of RNA substrate binding (*45*). This may explain why knock down of *Rnps1* or *Pnn* but not *Acin1* significantly reduced the intron detentions induced by the expression of mutant U2 (Figs. 6A and 6B). Our experiments did not support that SNIP1 reliably interact with the core EJC components (Figs. 5A and S6F). Given that ACIN1 and PNN are located in nuclear speckles - where post-transcriptional splicing occurs - it is plausible that RNPS1 regulates post-transcriptional splicing by temporarily forming the ASAP or PSAP complex, which is replaced by the core EJC complex during mRNA export (*70, 72–75*).

RNPS1 docking at DIs is sufficient to induce intron detention but has little effect on constitutive splicing (Figs. 6J and 6K). In addition, the interaction between SNIP1 and RNPS1 is required for spliceosome pausing at DIs (Figs. 6C-6J). We propose that SNIP1 and RNPS1 function a molecular brake for the spliceosome pausing (Fig. S10). That is why partial loss-of-function of SNIP1 or RNPS1, probably through releasing the molecular brake, rescues intron detentions caused by mutant U2. SNIP1 contains an FHA domain, which is involved in phospho-dependent protein/protein interaction (*43, 44*), and the FHA domain is required for the interaction between SNIP1 and RNPS1 (Fig. 6C). Therefore, phosphorylation/dephosphorylation status of RNPS1, especially in its S domain, may participates in the spliceosome pausing/resuming at highly regulated DIs. In fact, RNPS1 phosphorylation close to its S domain was proposed to regulate splicing *in vitro* and inhibition of SR protein kinases (Clk family members) were reported to promote the post- transcriptional splicing of DIs (*15, 20, 76*).

Like the *NMF291* mutation, *Snip1* KO decreases the splicing efficiency of DIs but has little effect on that of constitutively-spliced introns (Figs. 3A-3C and Fig. 7). These indicate that splicing of constitutively-spliced introns and DIs has different kinetics. Most likely, constitutive introns are co-transcriptional spliced but DIs undergo post-transcriptional splicing (*72*) (Fig. S10). In yeast, RES complex is not required for B complex formation but for the efficient transformation from B to B^act^ (*77*). Here, we suggest that SNIP1 may have two sides in regulation of DI splicing: 1) to form molecular brake together with RNPS1 to pause the spliceosome at DIs; 2) to facilitate the spliceosome transformation at DIs. Due to different splicing kinetics, DI but not constitutive intron splicing is primarily affected in presence of mutant (the *NMF291*^-/-^) or dysfunction (*Snip1*^-/-^) spliceosome (Fig. S10). Partial loss of SNIP1 or RNPS1 rescues mutant-U2 caused intron detentions probably through releasing the molecular brake, while complete removal of *Snip1* may impair the spliceosome transformation at DIs, thus lower the splicing efficiency (Fig. S10). Global accumulation of DIs will sequester spliceosome pausing complex, which in turn lowers the available spliceosome abundance and worsens DI splicing. Indeed, decreased amounts of core and non-core spliceosome components leads to accumulation of intron-containing transcripts (*78*). In addition, accumulation of intron-containing transcripts has been documented in patient brains with neurodegenerative diseases (*79, 80*). Given that genes involved in RNA metabolism/processing and cellular response to stress tend to transcribe IDTs, global accumulation of DIs will in turn further damages these gene functions. The mechanisms underlying post-transcriptional DI splicing we describe here may help to improve the understanding of pathogenesis of neurodegenerative diseases and to develop intervention strategies in the future.

## Materials and methods

### Mice

All the mice described in this work were maintained on the C57BL/6J background (JAX, Stock No. 000664) and housed in Tsinghua Animal Facility that has been accredited by the Association for the Assessment and Accreditation of Laboratory Animal Care International (AAALAC) since 2014. All animal protocols were approved by the Institutional Animal Care and Use Committee (IACUC) at Tsinghua University based on Guide for the Care and Use of Laboratory Animals (Eighth Edition, NHR). Mice were housed in isolated ventilated cages (maximum six mice per cage), on a 12/12-hour light/dark cycle, 22-26°C, 40–70% humidity with sterile pellet food and water *ad libitum*. Cages were checked daily to ensure animal welfare. Body weight was assessed regularly to ensure no weight loss. Whenever animals were used for research, we followed the 3Rs (replacement, refinement or reduction) rules. The C57BL/6J and ICR mice were purchased from Charles River Laboratories, Beijing, China. The Nse-CreER^T2^ line was imported from The Jackson Laboratory (JAX, Stock No. 022763) (*55*). For tamoxifen injection, we followed previous reports (*81, 82*). In brief, tamoxifen base (T5648-Sigma) was dissolved in corn oil and prepared freshly before use.

### ENU mutagenesis and modifier identification

For ENU-induced mutagenesis, the *NMF291*^-/+^ males were i.p. injected with ENU (80-110 mg/kg B. W.) for three consecutive weeks, as previously reported (*83, 84*). After an infertility test, the ENU-treated males were crossed to untreated *NMF291*^-/+^ females for G1. Of these offsprings, the *NMF291*^-/-^ mice were used for behavioral tests, and the mice with less ataxia phenotype and improved lifespan were selected for family pedigree determination.

For identification of the modifier candidates, the ENU family members with or without phenotypic improvement were applied for exome sequencing. For exome capture and library construction, the instructions of NimbleGen SeqCap EZ Exome Library SR Platform (Roche) were followed. Briefly, genomic DNA was fragmented to 200-300 bps with ultrasonic shearing. End-repair, A-tailing, adapter ligation, and pre-capture ligation were performed by using the KAPA LTP Library Preparation Kit (Roche). After exome capture, the resulting samples were amplified by KAPA HiFi HotStart Ready Mix using Pre-LM-PCR Oligos 1 & 2. Libraries passing QC were sequenced (Illumina HiSeq 2500 platform) and data processing and variant discovery were performed by using GATK platform (*85*). Variant annotation and classification were achieved using ANNOVAR (*86*).

### Generation of *Snip1* KO, Flag-tag KI, and conditional KO mice

The CRISPR design website (http://crispr.mit.edu) was used to design gRNAs and avoid off-targets (*87, 88*). Cas9 mRNA, gRNA (s), and/or donor DNA were injected to C57BL/6J embryos. Then the injected embryos were transferred to the oviduct ampulla of pseudo-pregnant ICR female mice. For generation of the *Snip1* KO mouse, the gRNA target sequence was: AGTGAGCGAGACCGGCACCG<u>GGG</u> (with PAM site underlined). For generation of the *Snip1* Flag-tagged mouse, the gRNA target sequence was: GGGACGGTTTCTAACAGTAG<u>AGG</u>. Donor DNA had two homology arms (∼200bps each) flanking the mutant PAM site. For generation of the *Snip1* floxed mouse, we employed multiple gRNAs to increase homologous recombination. The gRNAs were: gRNA-A1: GAGTCTAACTGGCCCTTCGG<u>GGG</u>, gRNA-A2: CAATGGTACCATCCTTTAAC<u>AGG</u>, gRNA-B1: AGTGTGGTTCTTCCCCCGAA<u>GGG</u>, gRNA-B2: CTGTTAAAGGATGGTACCAT<u>TGG</u>. Two LoxP sites were placed in the *Snip1* intron2 and intron3, respectively, away from conserved intronic sequences. The donor DNA contained the targeted exon3, the flanking two LoxP sites, and two homology arms (∼800bps each). C57BL/6J mouse genomic DNA was used as a template to amplify the sequences and the homology arms in the donor DNAs. Mice with the right genotypes were further crossed to C57BL/6J mice for at least three generations to establish the lines.

### Hematoxylin and eosin staining

Mice were anesthetized and transcardially perfused with PBS and then Bouin’s solution (Sigma-Aldrich). After post-fix in Bouin’s solution, the brain tissues were paraffin embedded. The paraffin sections were applied for hematoxylin and eosin (H&E) staining. The stained sections were scanned with Pannoramic Digital Slide Scanners (3DHISTECH).

### Cell culture, plasmid construction, transfection, lentiviral infection, shRNA knockdown, and NMD reporter assay

HEK293FT and Neuro-2a cells were cultured in Dulbecco’s modified Eagle’s medium (DMEM, Corning) complemented with 10% FBS, 1% penicillin–streptomycin in 5% CO2 at 37℃. For plasmid construction, genes were cloned into pCMV-3tag-4A and pCMV-3tag-1A (Agilent Technologies). In addition, we replaced EGFP and Cas9 from pLJ-EGFP and LentiCas9-Blast (Addgene) with genes of interest. Cell transfection was performed using Lipofectamine 3000 (Thermo Fisher Scientific). shRNA clones were purchased from lentiviral vector-based shRNA libraries (MISSION shRNA library, Sigma-Aldrich). Lentivirus packaging was performed as previous reported (*89*). The NMD reporter assay in N2a cells was performed as previous described (*90*). Briefly, N2a cells were transfected with pCI-Renilla/β-globin and pCI-firefly reporters (gifted from Drs. Gabriele Neu-Yilik and Andreas E. Kulozik). After 24 hours, cells were treated with100 μg/ml CHX or 0.01% DMSO for 5 hours. Renilla and firefly luciferase were detected using Dual-Luciferase Reporter System (Promega).

### Antibodies and immunofluorescence

Antibodies used in this study were: Mouse anti-FLAG (Abmart, M2008H), rabbit anti-FLAG (Sigma, F7425), mouse anti-TUBULIN (Sigma, T6793), mouse anti-HA (Abcam, ab130275), rabbit anti-HA (Invitrogen, 715500), mouse anti-Myc (Abmart, M20002), rabbit anti-NeuN (Cell Signaling, 24307), anti-c-Myc Magnetic Beads (Thermo Fisher Scientific, 88842), donkey anti-mouse IgG secondary antibody (Alexa Fluor 555, Thermo Fisher Scientific, A31570), donkey anti-rabbit IgG secondary antibody (Alexa Fluor 555, Thermo Fisher Scientific, A31572), donkey anti-goat IgG secondary antibody (Alexa Fluor 488, Thermo Fisher Scientific, A11055). PTBP1 N-terminal antibody (PTB-NT, generated from 1-15 a.a.) was gifted from Dr. Douglas Black.

For tissue immunostaining, mice were anesthetized and transcardially perfused with PBS and then 4% paraformaldehyde. Brain tissues were submerged in 10%, 20%, and 30% sucrose solution for gradient hydration. The resulting tissues were embedded in OCT and cut on a cryostat. The brain sections were blocked in 3% BSA and then incubated with primary antibody overnight. After 0.5% PBS-T washes, the sections were incubated with Alexa Fluor-conjugated secondary antibodies (Thermo Fisher Scientific). For cultured cell staining, cells were fixed with 4% paraformaldehyde and blocked with blocking buffer. The fixed cells were incubated with primary antibodies overnight. Images were taken by Nikon A1 confocal microscopy.

### Immunoblot, IP, and co-IP/MS

For immunoblot, fresh tissues were homogenized on ice with RIPA buffer (25 mM Tris-HCl pH 7.6, 150 mM NaCl, 1% NP-40, 1% sodium deoxycholate, 0.1% SDS) complemented with protease inhibitor cocktail (Roche) and phosphatase inhibitor cocktail (PhosSTOP, Roche). For cell lysate, the cultured cells were washed with PBS first and then lysed with RIPA buffer. The resulting lysates were centrifuged at 12,000 × g for 10 minutes. Supernatant was boiled with 2 × SDS loading buffer for immunoblot loading.

For IP, mouse cerebellum was homogenized on ice with IP buffer (150 mM KCl, 25 mM Tris pH 7.4, 5 mM EDTA, 1% Nonidet P-40) complemented with protease and phosphatase inhibitor cocktail (Roche). Cultured cells were washed with ice-cold PBS three times and lysed with IP buffer on ice. Tissue and cell lysates were rotated at 4℃ for 0.5 hour and then centrifuged to remove cell debris. The supernatant was collected and incubated with anti-FLAG M2 magnetic beads (M8823, Merck) 4℃ overnight. The magnetic beads were washed with IP buffer twice and TBS buffer (50 mM Tris-HCl, 150 mM NaCl, pH 7.4) twice. The resulting samples were boiled with 2 × SDS loading buffer.

For MS, samples were loaded on SDS-PAGE gels for separation, stained with SimplyBlue™ SafeStain (Life technologies), and excised. The gel slices were reduced and in-gel digested with sequencing grade modified trypsin (Promega). The resulting samples were quenched by 10% trifluoroacetic acid and peptides were extracted and dissolved in 0.1% trifluoroacetic acid. For LC-MS/MS, the purified peptides were separated by a C18 column (75 μm inner diameter, 150 mm length, 5 μm, 300 Å) and directly connected with an Orbitrap Fusion Lumos mass spectrometer (Thermo Fisher Scientific). Mobile phase A was an aqueous solution of 0.1% formic acid, and mobile phase B was 0.1% formic acid in acetonitrile. The MS/MS spectra were searched against the Uniport mouse database using Proteome Discoverer (version PD1.4, Thermo Scientific™). The peptide spectrum match (PSM) was calculated by Percolator provided by Proteome Discoverer, and only peptide FDR less than 0.01 was included for further analysis. Peptides only assigned to a given protein group were considered as unique. FDR was also set 0.01 for protein identifications.

### Proteome analysis for potential peptides encoded by DIs and their upstream exons

For proteome analysis, cerebellum was quickly removed and placed in a Dounce homogenizer. Tissues were lysed with 8 M urea in PBS supplemented with protease and phosphatase inhibitor cocktail (Roche) at room temperature. Lysates were centrifuged at 12,000 rpm for 10 minutes and supernatant was collected. Protein concentration was determined by BCA Protein Assay Kit (Pierce™). The tryptic peptides were fractionated with a XBridgeTM BEH300 C18 column (Waters, MA). LC-MS/MS analysis was similar to what we described above. To identify potential peptides encoded by DIs, a customized peptide database was generated as previously reported (*29*). Briefly, all DI events were extracted from wildtype and *NMF291*^-/-^ RNA-Seq results. The RNA sequences were in silico translated to peptides from 60 nucleotides upstream of DIs, until a PTC is encountered. The MS/MS spectra from each LC-MS/MS run were searched against the customized peptide database by the Sequest HT search engine of Proteome Discoverer, to identify potential peptides encoded by intronic sequences. Trypsin was specified as the proteolytic enzyme, and we allowed up to two missed cleavage sites. PSM was validated using Percolator, and only FDR < 0.01 was considered correct.

### Total RNA extraction, reverse transcription, real-time qPCR, and nuclear and cytoplasmic RNA extraction

Total RNA was extracted with TRIzol reagent (Thermo Fisher Scientific). The extracted RNA was dissolved in nuclease-free water and then treated with DNase (RQ1 RNase-Free DNase, Promega). Reverse transcription was achieved with M-MLV Reverse Transcriptase (Promega). Real-time qPCR was achieved with qPCR SYBR Green Mix (Yeasen).

For nuclear and cytoplasmic RNA extraction (*91*), N2a cells were placed on ice, washed with ice-cold PBS three times, and then collected by spinning at 1,000 rpm for 10 minutes. The resulting cells were resuspended in 200 μl lysis buffer A (pH 8.0 10 mM Tris, 140 mM NaCl, 1.5 mM MgCl2, 0.5% NP-40, 1 mM DTT, 100 U/ml RNasin), incubated on ice for 5 minutes, then centrifuged at 1,000 × g for three minutes. The supernatant was collected for cytoplasmic RNA extraction. The pellets were further washed twice with the lysis buffer A and once with the lysis buffer A complemented with 1% Tween-40 and 0.5% deoxycholic acid. The resulting pellets were resuspended in TRIzol for nuclear extraction.

### RNA-Seq

Total RNA was dissolved in nuclease-free water, treated with DNase (Ambion, Thermo Fisher Scientific), and quantified by Qubit RNA Assay Kit (Thermo Fisher Scientific). Agilent 2100 Bioanalyzer (Agilent Technologies) was employed to check the quality of RNA. Total RNA (3 μg) was applied for poly(A) mRNA purification by using oligo-d(T) magnetic beads (S1419S, NEB). RNA fragmentation, cDNA synthesis, terminal repair, A-tailing, and adapter ligation were performed by using RNA library prep kit (E7530L, NEB). DNA products were cleaned using AMPure XP beads (Beckman). Library quality was checked by Agilent 2100 Bioanalyzer and quantified by real-time PCR. Sequencing was performed on the Illumina HiSeq platform.

### IR analysis

The quality of RNA-Seq reads was assessed by FastQC. The adaptors and low-quality reads were removed by Cutadapt to obtain clean reads. The resulting reads were aligned to mouse genome mm10 using HISAT2 (*92*). IR was analyzed as previously reported with slight modifications (*28–30*). In brief, every intron in the genome was considered as a potential retained intron, while only introns with both flanking exons covered by the aligned reads were included for further analysis. To avoid the influence of small non-coding RNAs and unknown exons located inside of the introns and low mappability regions in large introns, including high GC regions and repetitive sequence, only reads covering exon-intron junctions and exon-exon junctions were used to calculate the IRI (intron retention index). IRI for 5’ SS and 3’ SS were calculated separately. We considered the exon-intron junction reads as intronic reads, and the sum of exon-intron and exon-exon junction reads as total reads. IRI = (5’SS intronic reads / 5’SS total reads + 3’SS intronic reads / 3’SS total reads)/2. For RNA-Seq data, IRI > 0.1 and intron coverage > 0.9 were set to identify reliable IR events. For the comparison of differential intron usage, DEXSeq was employed for the statistical analysis, which offers reliable control of *p*adj by estimation of dispersion for each counting bin with generalized linear models (*93*).

### Heatmap, GO analysis, and conservation analysis

Heatmaps of Z-scores were plotted using Heatmap.2 of the R package “gplots”. To compare reliable DI events across different conditions, only events with IRI > 0.1 were included in the analysis. For GO enrichment analysis, the PANTHER Classification System was used to test the overrepresentation of genes (*94*). Statistical overrepresentation testing was performed for the GO biological process complete category and the enriched GO terms were ranked by fold-enrichment and FDR and the redundant GO terms were manually removed. To determine whether DIs are conserved between human and mouse, intron coordinate conversions were performed using UCSC LiftOver (from mm10 to hg38). The highly conserved introns (Liftover remap ratio > 90% between human and mouse) were included for the conservation comparison (*p* values were calculated with two-sided proportion tests).

### Nanopore full-length mRNA sequencing

Mouse cerebellar total RNA was extracted using TRIzol and treated with DNase. RNA quality was assessed using an Agilent 2100 Bioanalyzer to ensure the integrity of RNA. Total RNA (1 μg) was used for cDNA library construction, following the protocols suggested by Oxford Nanopore Technologies (ONT). Reverse transcription and strand-switching were performed using a cDNA-PCR Sequencing Kit (SQK-PCS109, ONT). cDNA was PCR amplified for 14 cycles by using LongAmp Taq (NEB). ONT adaptor ligation was performed using T4 DNA ligase (NEB). The resulting DNA was purified by Agencourt XP beads. The final libraries were added to FLO-MIN109 flowcells (ONT), sequenced using PromethION platform. Raw ONT reads were filtered by read quality score (> 7) and minimal reads length (> 500bps). Clean ONT reads were aligned to mouse genome reference mm10. Aligned reads were converted to bam files, sorted, and indexed by SAMtools (*95*). Reliable full-length transcripts were filtered to contain 5’UTR and 3’UTR regions by using BEDTools (*96*). To identify reliable DIs, minimal IRI was set as 0.05 with intronic region coverage larger than 0.9.

### Native RIP-seq

Native RIP-seq was performed as previously reported with minor modifications (*97*). For mouse tissues, mouse cerebellum was removed and placed in a Dounce homogenizer on ice. Lysis was in RIP buffer (150 mM KCl, 25 mM Tris pH 7.4, 5 mM EDTA, 0.5 mM DTT, 0.5% Nonidet P-40) complemented with protease and phosphatase inhibitor cocktail (Roche) and 100 U/ml RNasin (Promega). For cultured cells, cells were washed with ice-cold PBS three times and lysed in RIP buffer on ice. Tissue and cell lysates went through a 23-gauge needle, rotated at 4℃ for 0.5 hour, and centrifuged to remove cell debris. The resulting supernatant was incubated with anti-FLAG M2 magnetic beads (M8823, Merck). After incubation for 12 hours at 4℃, M2 beads were washed with RIP buffer twice and TBS buffer twice. Elution was achieved by using 250 μg/ml 3 × Flag peptides (F4799, Sigma). The resulting eluates were resuspended in TRIzol for the following RNA extraction. Smart-seq2 was performed following a standard protocol as previously described (*98*). The amplified DNA (40 ng) was fragmented into ∼350 bp fragments using a Bioruptor® Sonication System (Diagenode Inc.). Illumina library construction was performed and qualified libraries were sequenced on the Illumina HiSeq platform. As described above, we used exon-intron junction and exon-exon junction reads to calculate IRI. For ratio comparison of IRI, IRI > 0.1 and intron coverage > 0.8 were set to identify reliable IR events.

### CLIP analysis

The RNPS1-GFP iCLIP and SF3a60 eCLIP were performed by Hauer et al. and Van Nostrand et al., previously (*52–54*), and the raw data were downloaded from ArrayExpress and ENCODE (*99–101*). The CLIP and corresponding RNA-Seq reads were aligned to the human reference genome hg19 by STAR v2.7 (*102*). The second read (R2) in each eCLIP read pair were extracted by SAMtools for downstream analysis, as described in the eCLIP-seq processing pipeline (*103*). To generate a CLIP read density plot, biological replicates were merged for combined reads count. Introns were grouped based on IR analysis in CLIP-related RNA-Seq data. DIs with high confidence were filtered by IRI > 0.1 and intron region coverage > 0.95, while efficiently spliced introns were filtered by IRI < 0.05. CLIP read coverage of introns and intron flanking exons were calculated using BEDTools (*96*), then read coverage were summed and normalized according to their relative position to 5’ SS and 3’ SS. The final read density was plotted using R (version 3.4.3). CLIP peak calling was achieved with CLIPper (CLIP peak enrichment recognition) by the default parameters (*104*). Binding stringency was ranked by the *p*-value calculated by CLIPper.

## ACKNOWLEDGMENTS

We thank Dr. Susan L. Ackerman for the *NMF291* mouse line. We thank the staff members in the Tsinghua Laboratory Animal Research Center (LARC) for mouse housing. We thank Dr. Haiteng Deng, Dr. Yi Huo, and Chongchong Zhao in the Tsinghua Proteomics Resource Center for mass spectrometry. We thank Dr. Douglas Black for the PTB-NT antibody and Drs. Gabriele Neu-Yilik and Andreas E. Kulozik for the NMD reporters. We also thank Dr. Michael Q. Zhang and Dr. Susan L. Ackerman for their comments on the manuscript. This work was supported by the National Natural Science Foundation of China (NSFC grants 31571097 and 81371361), the Thousand Talent Program for Young Outstanding Scientists, the Tsinghua-Peking Joint Center for Life Sciences, and IDG/McGovern Institute for Brain Research at Tsinghua.

## AUTHOR CONTRIBUTIONS

Y.J. and D.M. conceived the study and wrote the manuscript. D.M. performed the mouse forward genetic screening and identified the genetic modifier. L.L. performed the CRISPR-Cas9-mediated *Snip1* KO and *Snip1*-*Flag* KI in mice. D.M. performed the IR analysis and analyzed the RNA-Seq, nanopore sequencing, and CLIP data. X.Z. performed the immunostaining. D.M. performed the MS and RIP-seq and analyzed the data. D.M. and Q.Z. validated the RNA-Seq and MS results. D.M. and Q.Z. mapped the SNIP1/RNPS1 interaction regions.

## Supplemental information

**Figure S1.**
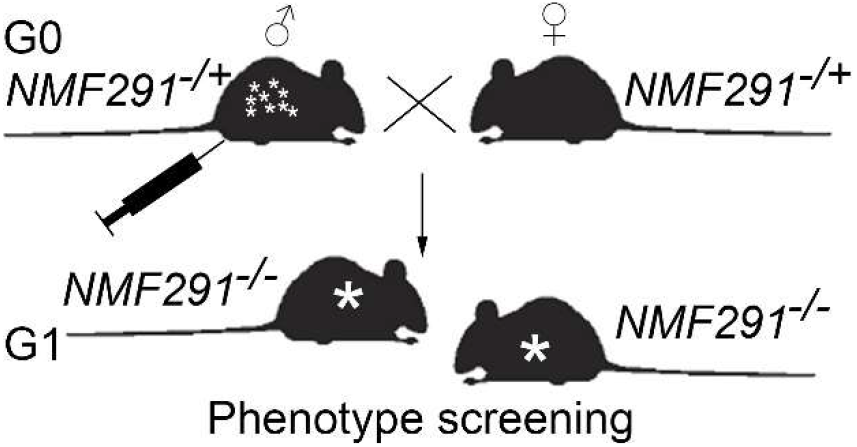
An ENU-induced mutagenesis screening for dominant modifier(s) that rescue(s) the *NMF291*^-/-^ phenotypes. G0 male *NMF291*^-/+^ mice were injected with ENU (N-ethyl-N-nitrosourea) once a week for 3 weeks. After their fertilities recovered, they were bred to ENU- untreated female *NMF291*^-/+^ mice. The G1 *NMF291*^-/-^ mice carrying less ataxia phenotype and improved lifespan were crossed to ENU-untreated *NMF291*^-/+^ mice to determine the family pedigree.

**Table S1.** Modifier candidates for the *NMF291* phenotypes.

| Gene | Position | SNV | Category | Rescue | No rescue |
| --- | --- | --- | --- | --- | --- |
| *Dnajc1* | chr2:18284715 | T/C | non-synonymous | 1/2 | 2/2 |
| *Fam193a* | chr5:34436543 | T/C | non-synonymous | 1/2 | 1/2 |
| *Ctr9* | chr7:111043186 | A/G | non-synonymous | 2/3 | 1/3 |
| *Gtf3c1* | chr7:125646516 | G/A | non-synonymous | 2/2 | 1/2 |
| *Nlrp1b* | chr11:71228365 | C/T | non-synonymous | 2/2 | 1/2 |
| *Pcdhac1* | chr18:37090163 | T/A | non-synonymous | 1/3 | 1/3 |
| *Ablim1* | chr19:57061311 | A/T | Stop-gain | 2/2 | 2/2 |
| *Frem1* | chr4:82914625 | T/A | Stop-gain | 6/7 | 2/5 |
| *Gm12794* | chr4:101941270 | T/G | non-synonymous | 18/18 | 1/14 |
| *Ptch2* | chr4:117114863 | T/A | 3'UTR | 8/8 | 1/9 |
| *Snip1* | chr4:125068193 | A/G | 5'SS/splicing | 27/27 | 0/19 |
| *Aunip* | chr4:134523450 | A/G | synonymous | 8/9 | 0/9 |
| *Kif17* | chr4:138291508 | T/A | non-synonymous | 7/8 | 0/10 |

Note: The modifier candidates were identified by an exome capture as previously described (PMID: 32051525). The ENU-induced *Snip1* mutation was co-segregated with the rescued phenotype among the candidates we examined, part of which are shown here. The mouse GRCm38/mm10 built was used as the reference genome. SNV, single nucleotide variant. Red, genotypes did not meet the expected phenotypes; green, genotypes agreed with the phenotypes.

**Table S2.** The abnormal Mendelian ratio seen in the progenies from *Snip1*^M+^ and *Snip1*^-/+^ intercrosses.

|  | *Snip1<sup>+/+</sup>* | *Snip1<sup>M/+</sup>* | *Snip1<sup>M/M</sup>* |
| --- | --- | --- | --- |
| Expected segregation ratio | 11 | 22 | 11 |
| Observed segregation ratio | 10 | 34 | 0 |

Note: The Chi-square value was equal to 17.64 ( $p < 0.001$ )

|  | *Snip1<sup>+/+</sup>* | *Snip1<sup>-/+</sup>* | *Snip1<sup>-/-</sup>* |
| --- | --- | --- | --- |
| Expected segregation ratio | 11.75 | 23.5 | 11.75 |
| Observed segregation ratio | 16 | 31 | 0 |

Note: The Chi-square value was equal to 15.68 ( $p < 0.001$ )

**Figure S2.**
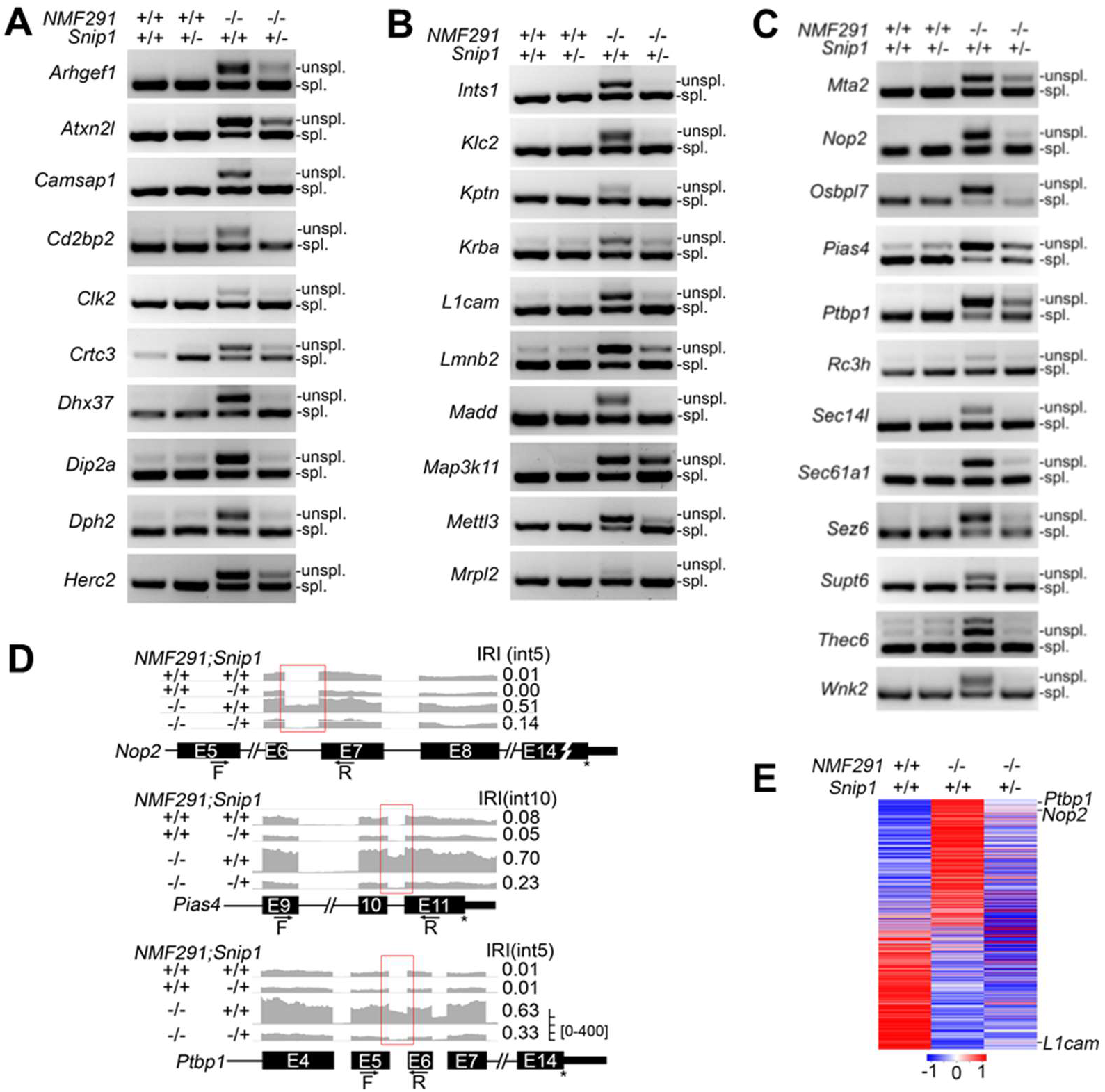
Haploinsufficiency of *Snip1* partially rescues the IRs and their corresponding gene expressions. (A-C) Validation of the IRs rescued by *Snip1*^-/+^ by RT-PCR. Cerebella were harvested at one month of age with the indicated genotypes. Unspl., unspliced transcripts; spl., spliced transcripts. (D) The representative IRs visualized by IGV. Primers used for the RT-PCR validation in (C) were illustrated. F, forward primer; R, reverse primer. (E) Corresponding genes (1455) of the rescued IRs (1961) shown in Figure 2C were included for expression analysis. The Z-score was used to normalize expression level in each row.

**Figure S3.**
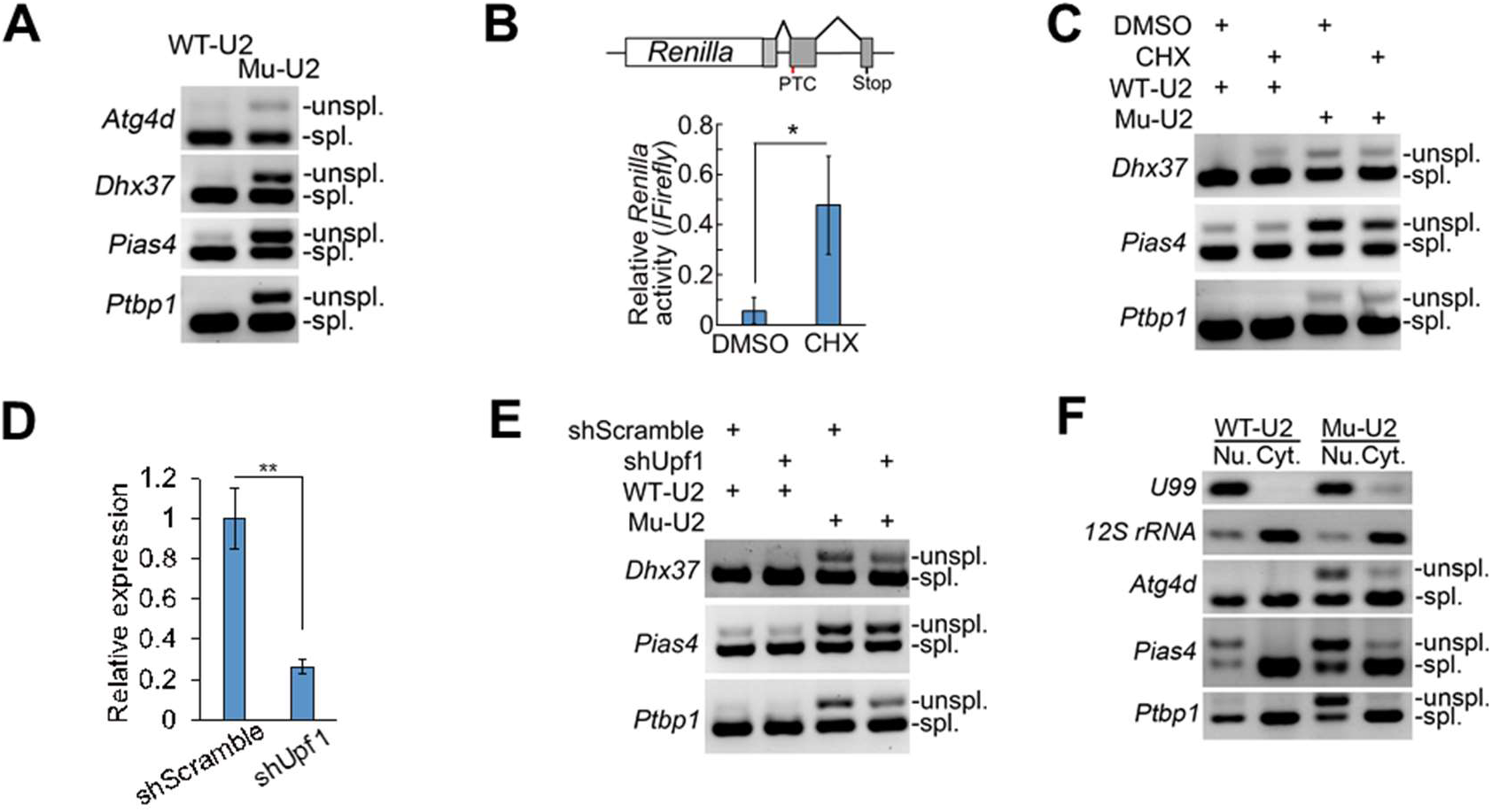
Intron-containing transcripts overrepresented in *NMF291* mutant cerebellum are likely IDTs. (A) The DIs amplified by Mu-U2 but not WT-U2 in N2a cells. (B) Previously described a NMD reporter (PMID: 1693475) in which *Renilla* luciferase contains a PTC in the second last exon. Application of CHX significantly inhibited NMD measured by relative *Renilla* activity. The *Firefly* luciferase was used to normalize transfection efficiency. (C) The intron-containing transcripts were insensitive to application of CHX in culture medium of N2a cells expressing WT-U2 or Mu-U2. (D and E) N2a cells were infected with *Upf1* and scrambled shRNA (MissionRNAi, Sigma). The relative expression level of *Upf1* measured by quantitative-PCR (D). The intron-containing transcripts were detected by RT-PCR (E). (F) The nuclear and cytosolic fractions evidenced by the enrichments of *U99* and *12s rRNA*, respectively. Nu., nucleus; Cyt., cytosol. In B and D, the values are presented as mean ± SEM, \**p*<0.05, \*\**p*<0.01, n = 4, t-test, SPSS. In A, C, E, and F, unspl., unspliced transcripts; spl., spliced transcripts.

**Figure S4.**
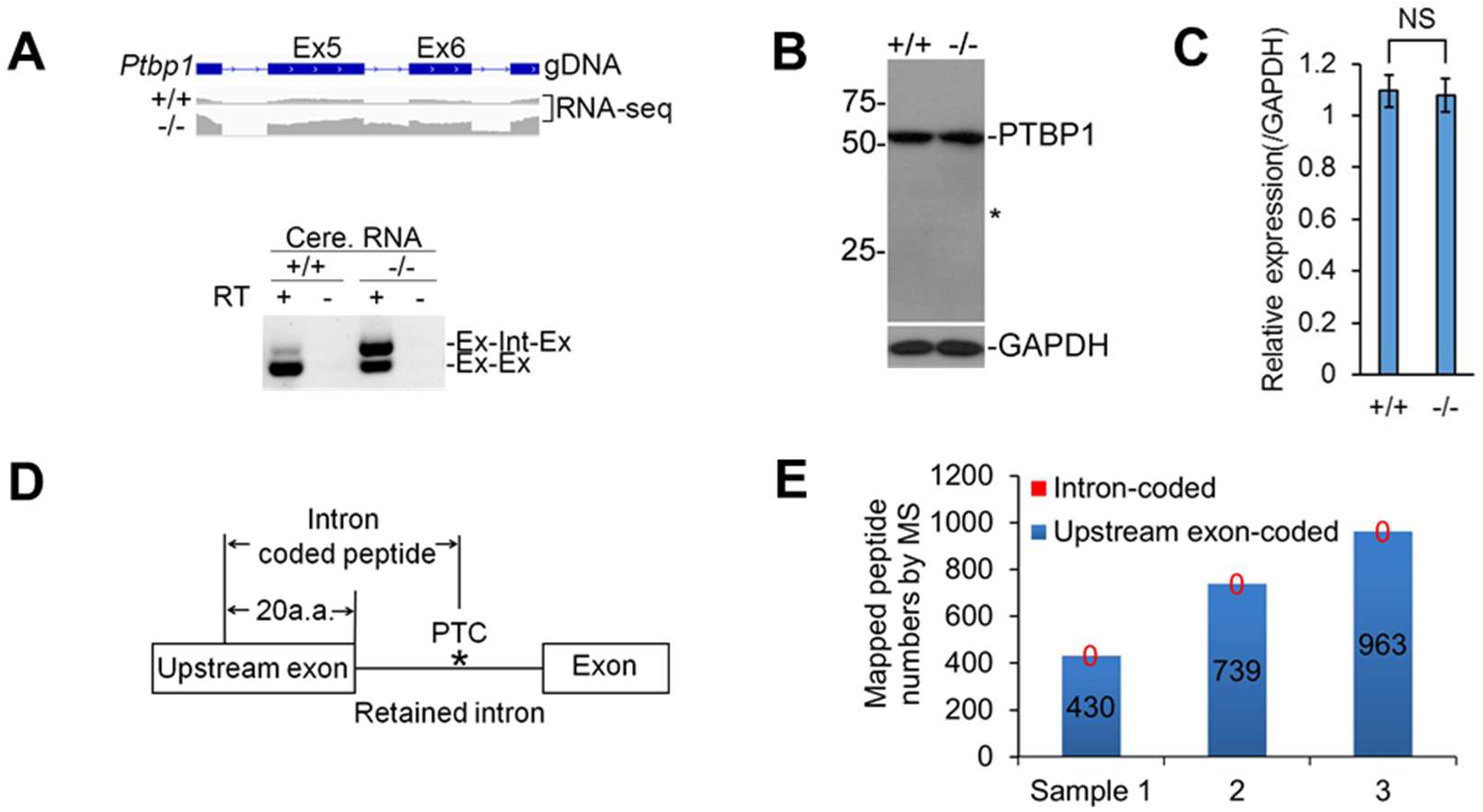
DIs are likely not code proteins. (A) Detained intron 5 of *Ptbp1* was evidenced by both RNA-seq (upper) and RT-PCR (lower) in wildtype (+/+) and *NMF291* mutant (-/-) cerebella. (B and C) The expression of PTBP1 by immunoblot. GAPDH served as loading control. Asterisk marked the corresponding molecular weight of PTBP1 supposedly coded by the intron 5-containing transcript. (D) A pipeline combines RNA-seq and MS (mass spectrometry) for seeking intron-coded peptide. Based on the DIs we identified in the *NMF291*^-/-^ cerebellum, we generated customized peptide database, which are coded by the DIs and their upstream exons. (E) Protein lysates from +/+ (n = 3) and *NMF291*^-/-^ mutant (n = 3) cerebella were applied for MS. The MS hits of our customized peptide database from one +/+ and mutant pair were pooled and summarized as one smaple. Mouse, one month of age. In A, unspl., unspliced transcripts; spl., spliced transcripts. In C, the values are presented as mean ± SEM. NS, no statistical significance (n = 3, t-test, SPSS).

**Figure S5.**
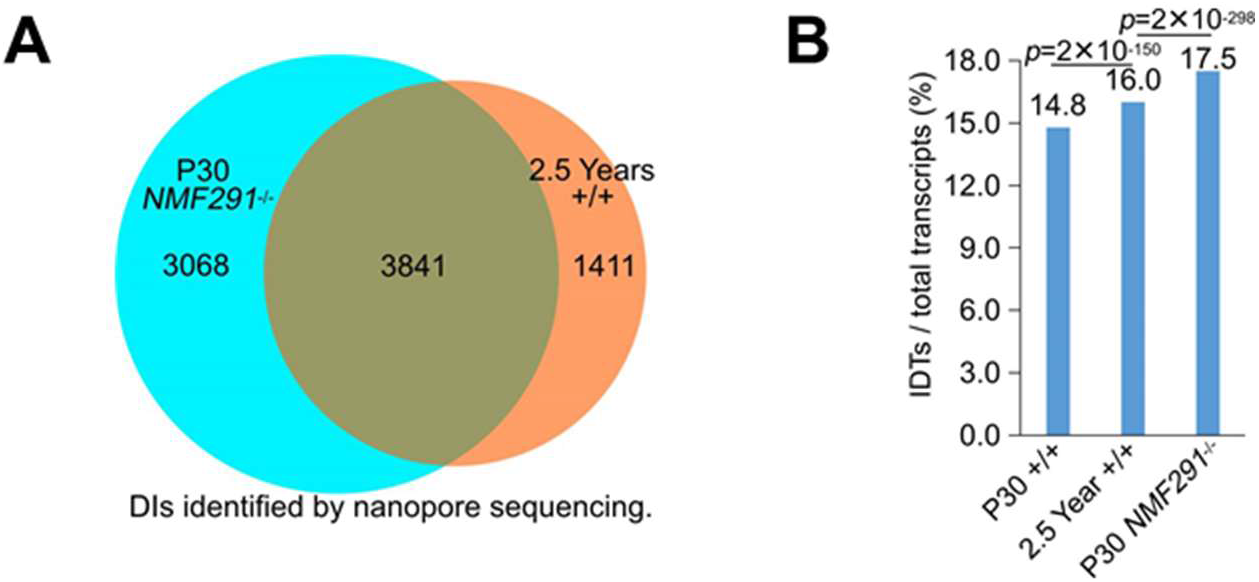
IDTs overrepresented in the *NMF291* mutant mouse are also accumulated in aged cerebellum, revealed by nanopore sequencing. (A) The majority of DIs accumulated in aged wildtype cerebellum are overlapped with that of the P30 *NMF291*^-/-^ cerebellum. (B) Percentage of IDTs in young (P30), aged (2.5 years), and P30 *NMF291*^-/-^ cerebella. All full-length nanopore reads containing 5’ UTR and 3’ UTR were included for the calculation. *p* values correspond to two-sided proportion tests.

**Figure S6.**
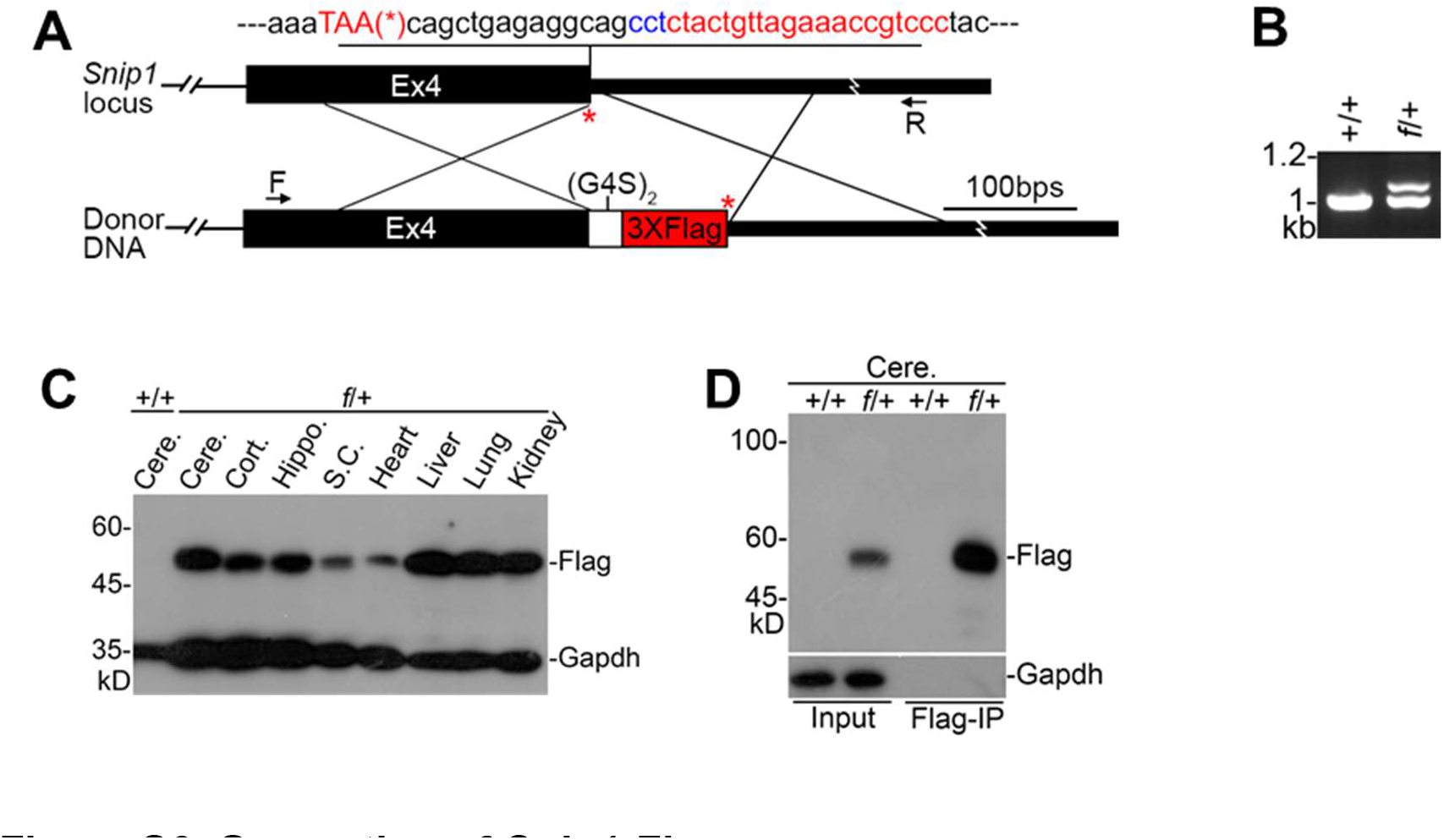
Generation of *Snip1-Flag* mouse. (A) Crispr/Cas9-based *Snip1-Flag* knockin (KI) design. The mouse *Snip1* locus and *Snip1* last coding exon (exon 4, Ex4) are shown (Upper). The stop codon (TAA) labeled with an asterisk (*); the target sequences of gRNA and PAM site (NGG) labeled in red and blue, respectively. In the donor DNA, a 3- time *Flag* tag was fused with *Snip1* last coding exon. A linker placed between Ex4 and the *Flag* sequences, which codes 2-time G4S (G4S)_2_. (B) Genomic DNA PCR confirmed the right KI. Primers for the PCR were labeled in (A) by arrows. F, forward; R, reverse. *f*/+, mouse heterozygous for *Snip1*-*Flag* KI. (C) SNIP1-Flag is ubiquitously expressed in various adult mouse tissues (mouse age, one month). (D) Flag-IP was performed with cerebellar protein lysates from *Snip1-Flag* KI mouse.

**Figure S7.**
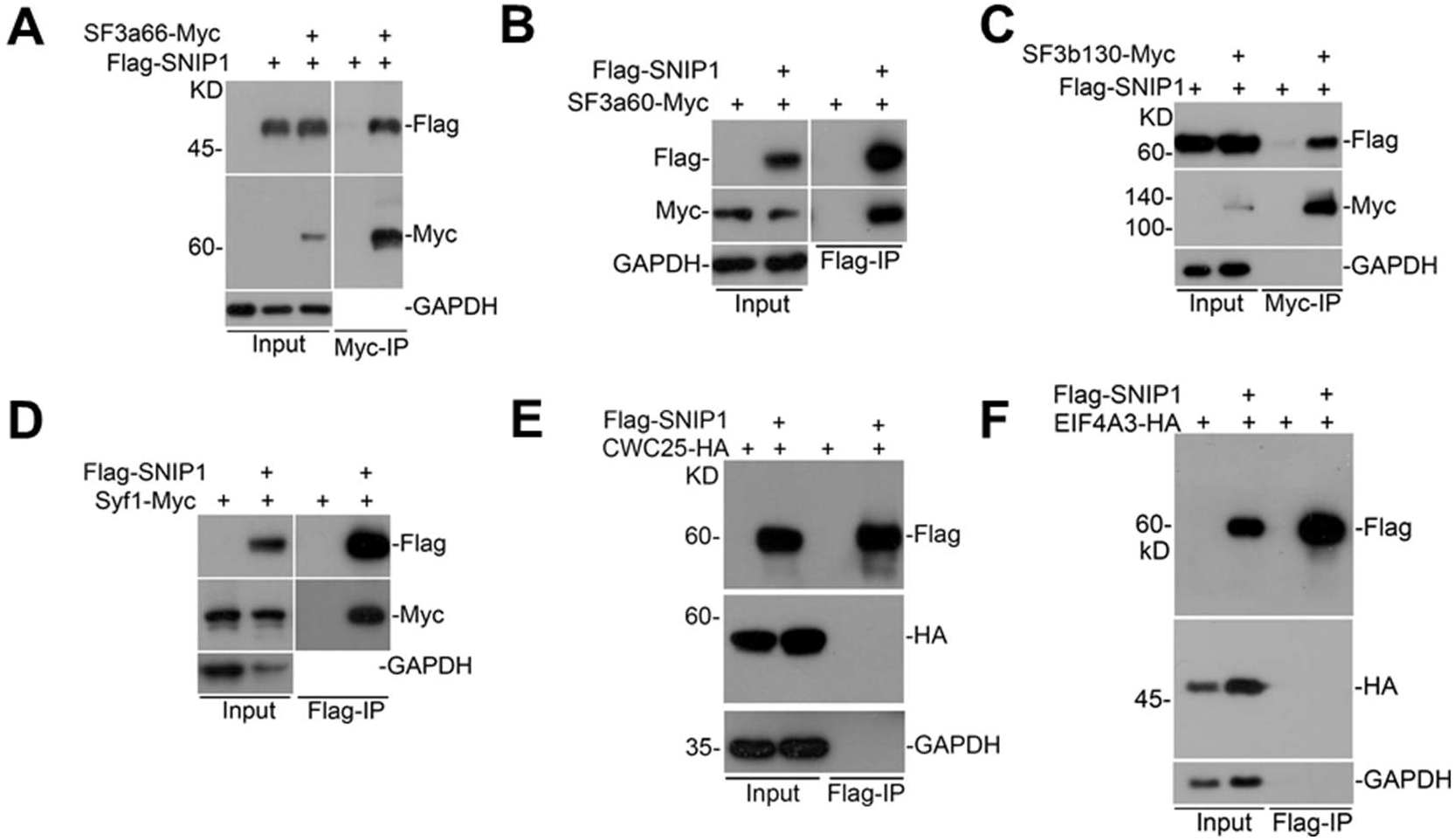
SNIP1-interacting partners are protein components found in B^act^ but not B*. (A-F) Interactions between SNIP1 and protein components found in B^act^ (SF3a66, SF3a60, SF3b130, and Syf1) but not those in B* (CWC25 and EIF4A3) were validated in N2a cells expressing the indicated tagged proteins.

**Figure S8.**
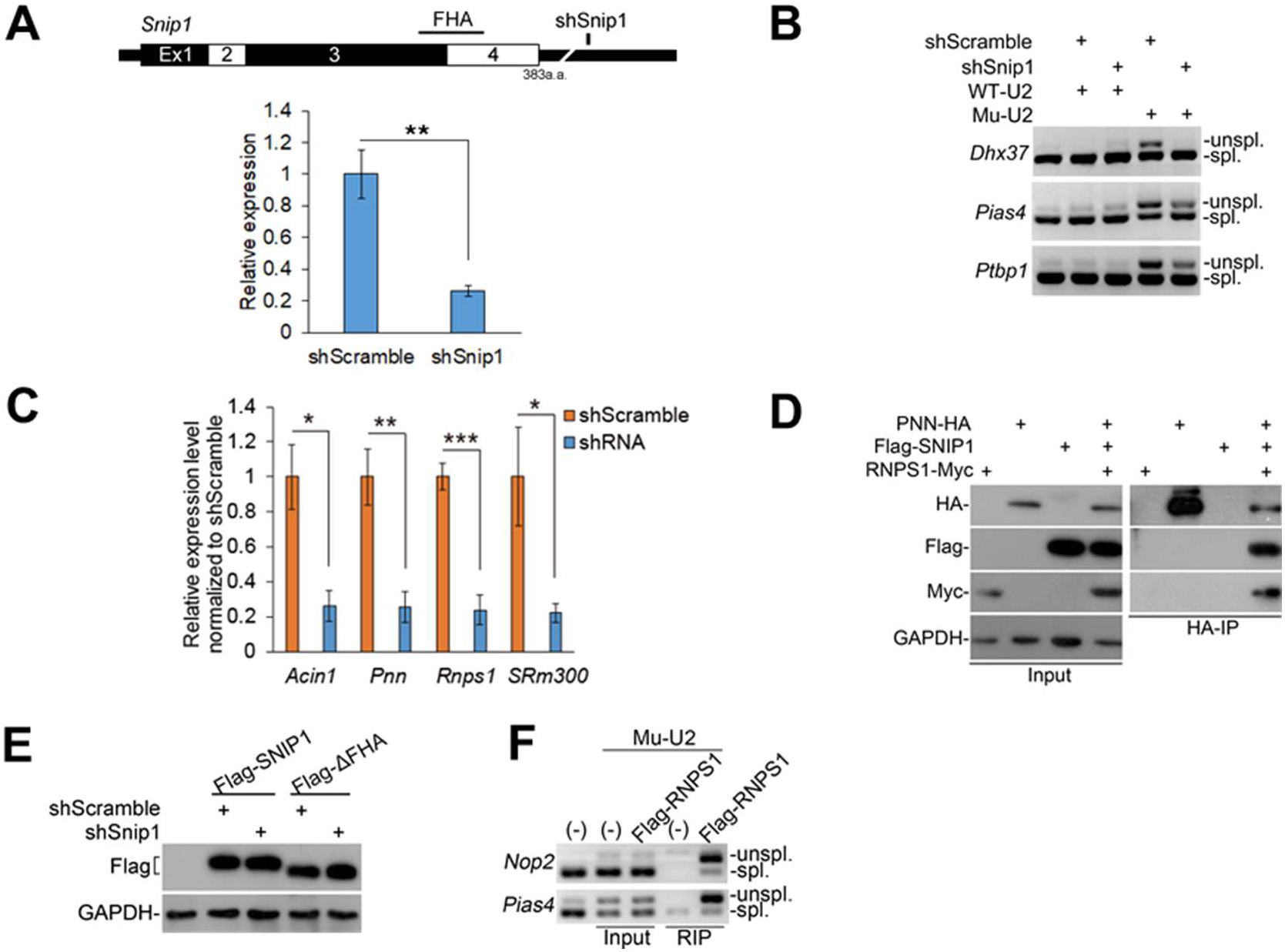
SNIP1 and RNPS1 function as a molecular brake to pause spliceosome at DIs. (A) Knockdown of *Snip1* in N2a cells by lentiviral shRNA (shSnip1). Note the target sequence of shSnip1 located in the 3’UTR (upper). The relative expression level of *Snip1* measured by quantitative-PCR (lower). The scrambled shRNA served as a control. (B) Knockdown of *Snip1* reduced DIs amplified by Mu-U2. The shRNA infected N2a cells were transfected with WT- and Mu-U2 expression plasmids, respectively. Scrambled shRNA infection served as a control. (C) Knockdown of genes encoding peripheral EJC components (*Acin1*, *Pnn*, and *Rnps1*) and a spicing factor (*SRm300*) measured by quantitative-PCR. (D) The interactions of SNIP1, PNN, and RNPS1 were evidenced by co-IP in N2a cells expressing PNN-HA, Flag-SNIP1, and RNPS1-Myc, simultaneously. (E) The shSnip1 targets 3’-UTR of endogenous *Snip1*, which does not affect the expression of exogenous full-length and ΔFHA SNIP1 in N2a cells. (F) Flag-RIP was performed with protein lysates from N2a cells infected with Flag-RNPS1 and transfected with Mu-U2 expression plasmid. In A and C, the values are presented as mean ± SEM, n = 4. \**p*<0.05, \*\**p*<0.01, \*\*\**p*<0.001, N.S., no statistical significance, t-test or ANOVA, SPSS. In B and F, DIs were measured by RT-PCR. Unspl., unspliced transcripts; spl., spliced transcripts.

**Figure S9.**
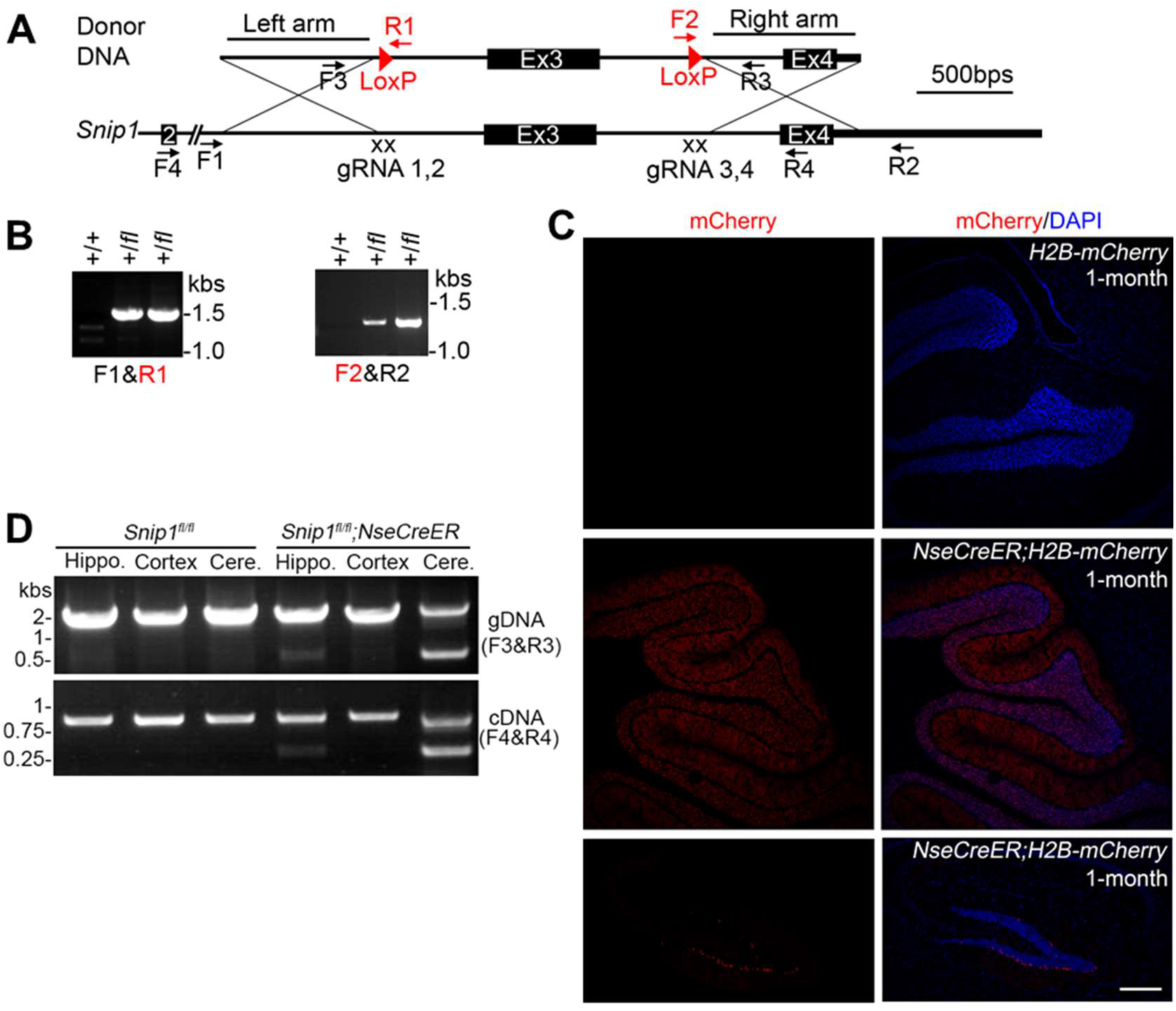
Generation of *Snip1* cKO mouse. (A) Crispr/Cas9-based *Snip1* cKO design. Genomic structure of mouse *Snip1* is illustrated. Homology arms (Left and Right) in donor DNA were labeled. Four gRNAs (2 gRNAs each side) flanking the *Snip1* exon3 (Ex3) were employed to increase homologous recombination. Primers for genotyping and detecting Cre-induced Ex3 deletion were labeled. Primers (R1 and F2) covering LoxP sites were labeled in red. F1 and R2 are located outside the homology arms. (B) Genomic PCR detected right LoxP site insertions in mice heterozygous for the floxed allele (+/*fl*). Primers used were labeled in (A). (C) Confirmation of the *NseCreER* expression by a previously described Cre-reporter, *H2B mCherry* (PMID: 25913859). The *NseCreER* were predominantly expressed in cerebellar granule cells and a few expressed in the granule neurons in hippocampal dentate gyrus. (D) Cre-dependent Ex3 deletion was detected at both genomic DNA (gDNA) and RNA levels in cerebellum and hippocampus but not in cortex in *Snip1^fl^*^/^*^fl^*;*NseCreER* mouse. *Snip1^fl^*^/^*^fl^* mouse served as negative control. Tamoxifen injections on P3, 4, and 5; DNA/RNA harvest on P7.

**Figure S10.**
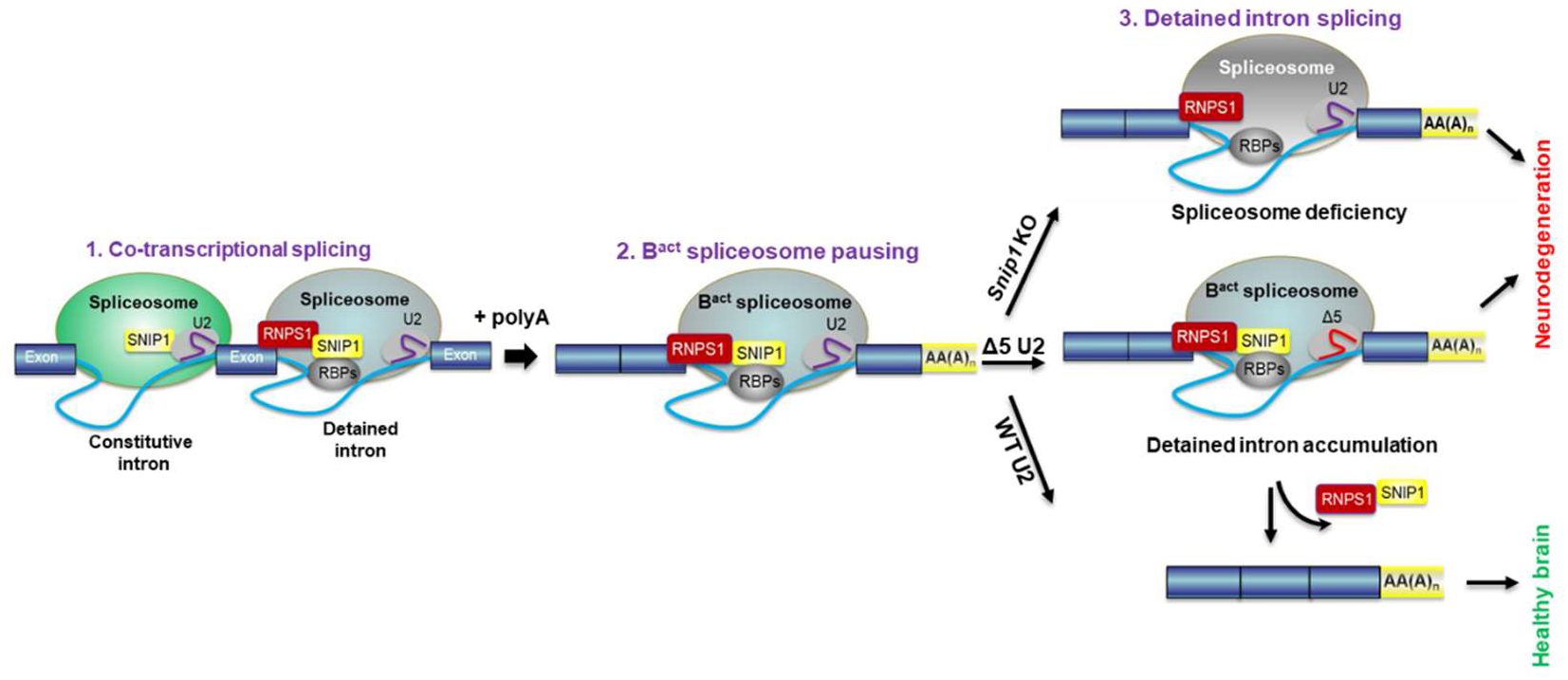
Our working model for post-transcriptional spliceosome pausing at highly regulated DIs and its contribution to neurodegeneration. We suggest that DI splicing is paused at B^act^ state, an active spliceosome but not catalytically primed, and SNIP1 and RNPS1 function as a molecular brake for the spliceosome pausing. The whole process can be divided into several steps. 1) Spliceosome loaded at both constitutive and detained introns. After constitutive introns are spliced, pre-mRNAs are polyadenylated to complete the co-transcriptional splicing. 2) The splicing of DIs is paused at B^act^ state (spliceosome pausing), which is mediated by SNIP1/RNPS1 containing complex. SNIP1, RNPS1 and B^act^ component preferentially dock at DIs, forming a molecular brake to modulate spliceosome pausing. RNPS1 recognizes the DIs and its neighboring sequences through its RRM (RNA recognition motif). Interaction between SNIP1 and RNPS1 is mediated by SNIP1 FHA (forkhead-associated) domain and RNPS1 S (serine-rich) domain. 3) The paused spliceosome is resumed, and DIs are spliced. The completely spliced transcripts are exported to cytosol for protein translation. Mutant (the *NMF291*^-/-^) or dysfunctional (*Snip1*^-/-^) spliceosome decreases splicing efficiency of DIs but has little effect on that of constitutively-spliced introns. That leads to accumulation of DIs, which in turn worsens DI splicing by sequester of spliceosome pausing complex and depletion of the available complex components and further damages the corresponding gene functions, and eventually causes neurodegeneration. Partial loss of SNIP1 or RNPS1 rescues intron detentions caused by expression of mutant U2, probably through releasing the molecular brake.

